# The validity of pairwise models in predicting community dynamics

**DOI:** 10.1101/060988

**Authors:** Babak Momeni, Li Xie, Wenying Shou

## Abstract

Pairwise models are commonly used to describe many-species communities. In these models, a focal species receives additive fitness effects from pairwise interactions with other species in the community (“pairwise additivity assumption”), and all pairwise interactions are represented by a single canonical equation form (“universality assumption”). Here, we analyze the validity of pairwise modeling. We build mechanistic reference models for chemical-mediated interactions in microbial communities, and attempt to derive corresponding pairwise models. Even when one species affects another via a single chemical mediator, different forms of pairwise models are appropriate for consumable versus reusable mediators, with the wrong model producing qualitatively wrong predictions. For multi-mediator interactions, a canonical model becomes even less tenable. These results, combined with potential violation of the pairwise additivity assumption in communities of more than two species, suggest that although pairwise modeling can be useful, we should examine its validity before employing it.

## Introduction

Multispecies microbial communities are ubiquitous. Microbial communities are important for industrial applications such as cheese and wine fermentation (van Hijum, Vaughan, and Vogel 2013) and municipal waste treatment (Seghezzo et al. 1998). Microbial communities are also important for human health: they can modulate immune responses and food digestion (Round and Mazmanian 2009; Kau et al. 2011) or cause diseases (Kelly 1980).

Community-level properties (e.g. species composition and biochemical activities) cannot be achieved, or achieved to the same extent, by summing the contributions of individual member species. Community-level properties are influenced by interactions wherein individuals alter the physiology of other individuals. To understand and predict properties of communities, choosing the appropriate mathematical model to describe species interactions is critical.

Two commonly-used modeling approaches are mechanistic modeling and pairwise modeling, each with its pros and cons. In mechanistic modeling, interaction mechanisms are explicitly modeled (Fig 1A and B, left panels). Thus, a mechanistic model requires discovering and quantifying interaction mechanisms (Fig 1 Table, “parameter” rows under “Mech.” column).

Such a mechanistic model can in principle quantitatively predict community dynamics when species evolution is negligible. However, the complexity of microbial interactions and the difficulty in identifying and quantifying interactions have made it challenging to construct mechanistic models.

In contrast to mechanistic modeling, pairwise modeling considers only the fitness effects of pairwise species interactions (Figs 1A and B, right panels). Pairwise models have two central assumptions. First, the “universality” assumption: Regardless of interaction mechanisms, how one species affects another can be abstracted into a single canonical equation form so that only parameters can vary among interactions. Second, the “pairwise additivity” assumption: a focal species receives additive fitness effects from pairwise interactions with other species in the community. Even though pairwise models do not capture the dynamics of chemical mediators, predicting species dynamics is still highly desirable in, for example, forecasting species diversity and compositional stability.

Pairwise models are easy to construct because they do not require knowledge of interaction mechanisms and need fewer parameters than mechanistic models (Fig 1 table). Parameters are relatively easy to estimate using community dynamics (Stein et al. 2013), or more systematically, using dynamics of monocultures and pairwise cocultures (Fig 2).

Not surprisingly, pairwise modeling has been commonly applied to communities (Wootton and Emmerson 2005). Pairwise models are often justified by their success in predicting ecological dynamics of two-species communities of prey-predation (Fig 1-FS1) (Volterra 1926; Wangersky 1978; “BiologyEOC - PopulationChanges” 2016) and competition (Gause 1934a; Gause 1934b). Pairwise modeling has been extended to model communities of more than two species (defined as *“multispecies communities”*),with empirical support from, for example, an artificial community of four competing protozoa species (Vandermeer 1969). Multispecies pairwise models have been extensively used to predict how perturbations to steady-state species composition exacerbate or decline over time (May 1972; Cohen and Newman 1984; Pimm 1982; Thébault and Fontaine 2010; Mougi and Kondoh 2012; Allesina and Tang 2012; Suweis et al. 2013; Coyte, Schluter, and Foster 2015).

However, pairwise modeling has known limitations. For instance, in a multispecies community, an interaction between two species can be altered by a third species (Levine 1976; Tilman 1987; Wootton 2002; Werner and Peacor 2003; Stanton 2003). Indirect interactions via a third species fall under two categories (Wootton 1993), which can be illustrated using the example of carnivore, herbivore, and plant. In an “interaction chain” (also known as “density-mediated indirect interactions”), a carnivore affects the density of an herbivore which in turn affects the density of plants. In “interaction modification” (also known as “trait-mediated indirect interactions” or “higher order interactions” (Vandermeer 1969; Wootton 1994; Billick and Case 1994; Wootton 2002)), a carnivore affects how often an herbivore forages plants. Interaction modification (but not interaction chain) violates the pairwise additivity assumption (Methods-Section 1). Interaction modification is thought to be common in ecological communities (Werner and Peacor 2003; Schmitz, Krivan, and Ovadia 2004). Limitation of pairwise modeling has also been studied experimentally (Dormann and Roxburgh 2005). However, empirically-observed failure of multispecies pairwise models could be due to limitations in data collection and analysis (Case and Bender 1981; Billick and Case 1994).

Given the benefits, limitations, and intellectual influence of pairwise modeling, we examine conditions under which pairwise models produce realistic predictions. Instead of investigating natural communities where interaction mechanisms can be difficult to identify, we start with *in silico* communities where species engage in predefined chemical interactions of the types commonly encountered in microbial communities. Based on these interactions, we construct mechanistic models, and attempt to derive from them pairwise models. A mechanistic reference model offers several advantages: community dynamics is deterministically known; deriving a pairwise model is not limited by inaccuracy of experimental tools; and the flexibility in creating different reference models allows us to explore a variety of conditions. This has allowed us to examine the domain of validity for pairwise modeling.

## Results

### Establishing a mechanistic reference model

In our mechanistic models (Fig 1A, left), we focus on chemical interactions which are widespread in microbial communities (Fig1-FS2) (Stams 1994; Czárán, Hoekstra, and Pagie 2002; Duan et al. 2009). A mechanistic model includes a set of species as well as chemicals that mediate interactions among species. Species **S_i_** could release or consume chemical **C_j_**, and chemical **C_j_** could increase or decrease the growth rate of species **S_k_**.

We assume that fitness effects from different chemical mediators on a focal species are additive. Not making this assumption will likely violate the additivity assumption essential to pairwise modeling. Additive fitness effects have been observed for certain “homologous” metabolites. For example, in multi-substrate carbon-limited chemostats of *E. coli*, the fitness effects from glucose and galactose were additive (Lendenmann and Egli 1998). “Heterologous” metabolites (e.g. carbon and nitrogen sources) likely affect cell fitness in a multiplicative fashion. However, if released mediators cause small changes to the concentrations already in the environment, then additivity approximation may still be acceptable. For example, suppose that the fitness influences of released carbon and nitrogen with respect to those already in the environment are *wc* and *wn*, respectively. If *w_c_*, *w_n_*<<1, the additional relative fitness influence will be (1+*w_c_*)(1+*w_n_*)-1 ≈*w_c_*+*w_n_*. However, even among homologous metabolites, fitness effects may not be additive (Hermsen et al. 2015). “Sequential” metabolites (e.g. diauxic shift) provide such examples.

We also assume that resources not involved in interactions are never limiting. We thus simulate continuous community growth similar to that in a turbidostat, diluting the total population to a low density once it has reached a high-density threshold. Within a dilution cycle, a mechanistic model can be represented by a set of first-order differential equations, as

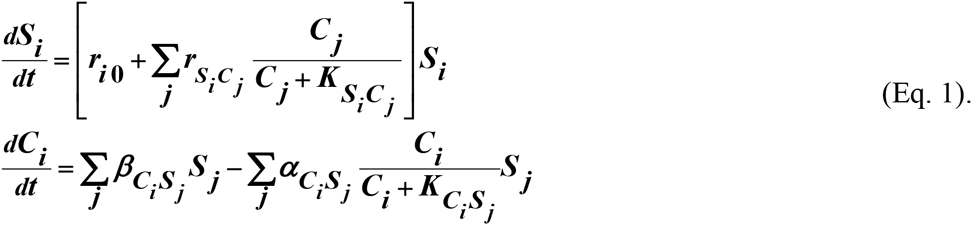

***S**_i_* and ***C**_i_* are state variables representing the concentrations of species **S_i_** and chemical **C_i_**, respectively. ***r_io_*** is the *basal fitness* of an individual of species **S_i_** (the net growth rate of a single individual in the absence of any intra-species or inter-species interactions). ***^r^S_i_C_j_*** reflects the maximal influence of chemical **C_j_** on the growth rate of **Si**, while ***^K^_S_i_C_i__*** is the concentration of **C_j_** achieving half maximal influence on the growth rate of **Si**. ***^β^C_i_S_i_***. and ***^α^C_i_S_j_*** are respectively the release rate and the maximum consumption rate of **Ci** by species **S_j_**. ***^K^C_i_S_j_*** is the ***C_i_*** at which half maximal rate of consumption by *S_j_* is achieved. All parameter definitions are summarized in Fig 1 table.

### Deriving a pairwise model

Ideally we would want a canonical pairwise model to represent the fitness effect of one species on another regardless of interaction mechanisms. Specifically, an N-species pairwise model is:

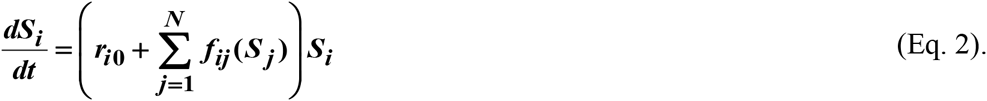

Here, ***r*_i0_** is the basal fitness of an individual of the focal species **S_i_ *f_ij_*(*S_j_*)** describes how ***S_j_***, the density of species **S_j_**, positively or negatively affects the fitness of **S_i_**. When ***j=i***, ***f_ii_***(***S_i_***) represents intra-population density-dependent fitness effect on **S_i_** (e.g. inhibition or stimulation of growth at high cell densities). The pairwise additivity assumption means that ***f_ij_***(***S_j_***) is a linear or nonlinear function of only ***S_j_*** and not of a third species.

***f_ij_*(*S_j_*)** can have several variations (Wangersky 1978): basic Lotka-Volterra, where the fitness influence of **S_j_** on **S_i_** linearly increases with the abundance of **S_j_** (Solé and Bascompte 2006); logistic Lotka-Volterra, which considers resource limitation by specifying a carrying capacity for each species (Thébault and Fontaine 2010; Mougi and Kondoh 2012); Lotka-Volterra with delayed influence, where the fitness influence of one species on another may lag in time (Gopalsamy 1992), and saturable Lotka-Volterra, where the fitness effect of **S_j_** on **S_i_** saturates at high density of **S_j_** (Thébault and Fontaine 2010). Since we model continuous growth which does not impose carrying capacity and since chemical influence from one species to another is likely saturable, we have adopted the saturable Lotka-Volterra as our canonical pairwise model:

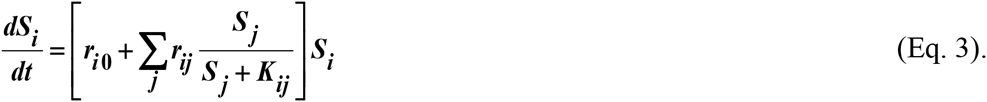

Here, ***r_ij_*** is the maximal positive or negative fitness effect of **S_j_** on **S_i_**, and ***K_ij_*** is the ***S_j_*** exerting half maximal fitness influence on **S_i_** (parameter definition in Fig 1 table). When ***j=i***, nonzero ***r_ii_*** and ***K_ii_*** reflect density-dependent growth effect in **S_i_**.

From a mechanistic model, we derive a pairwise model either analytically or numerically (Fig 2A). In the latter case (Fig 2B-C, Methods-Section 2), we should already have a pre-specified pairwise model (e.g. the canonical pairwise model) in mind. We then use the dynamics of monocultures and pairwise cocultures obtained from the mechanistic model to find parameters that minimize the difference between the two models within a training time window ***T***. Specifically, we define a distance measure 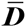 as the fold-difference between the dynamics from the two models, averaged over any time interval ***τ*** and species number ***N***(Fig 2C):

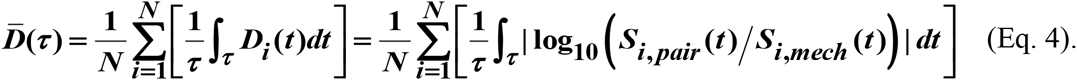

Here ***S**_i,pair_* and ***S**_i,mech_* are ***S**_i_* calculated using pairwise and mechanistic models, respectively. Since species with densities below a set extinction limit, ***S**_ext_*, are assumed to have gone extinct in the model, we set all densities below the extinction limit to ***S**_ext_* in calculating 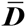 to avoid singularities. Within the training window **T**, minimizing 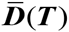 using a nonlinear least square routine yields parameters of the best-matching pairwise model. We then use 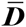 outside the training window to quantify how well the best-matching pairwise model predicts the mechanistic model.

### Reusable and consumable mediators are best represented by different forms of pairwise models

To build a pairwise model, we must accurately represent the fitness influence of one species on another (***r_ij_*** and ***K_ij_***). Even though this basic process seems straightforward as outlined in Fig 2, in practice, challenges may arise. For example, identifying the set of best-matching parameters for nonlinear functions may not be straightforward, and measurement errors further hamper parameter estimation. Partly due to these challenges, studies on deriving pairwise model parameters for a given community are scarce (Pascual and Kareiva 1996; Stein et al. 2013), despite the popularity of pairwise models. In this section, we analytically derive pairwise models from mechanistic models of two-species communities where one species affects the other species through a single mediator. The mediator is either reusable such as signaling molecules in quorum sensing (Duan et al. 2009; N.S. Jakubovics 2010) or consumable such as metabolites (Stams 1994; Freilich et al. 2011). To facilitate mathematical analysis, we consider community dynamics within a dilution cycle. We show that a single canonical pairwise model may not encompass these different interaction mechanisms.

Consider a commensal community where species **S_1_** stimulates the growth of species **S_2_** by producing a reusable (Fig 3A) or a consumable chemical **C_1_** (Fig 3B). When **C_1_** is reusable, the mechanistic model can be transformed into a pairwise model (Fig 3A), provided that the concentration of the mediator (which is initially zero) has acclimated to be proportional to the producer population size (Fig 3A; Fig 3-FS1). This pairwise model takes the canonical form (compare with Fig 1B right). Thus, the canonical pairwise model is appropriate, regardless of whether the producer coexists with the consumer, outcompetes the consumer, or is outcompeted by the consumer.

If **C_1_** is consumable, different scenarios are possible within a dilution cycle (Methods-Section 3).

Case I: When supplier **S_1_** always grows faster than consumer **S_2_** (***r*_10_ > *r*_20_ + *r_S_2_C_1__***), **C_1_** will accumulate within each dilution cycle (proportional to ***S*_1_**) without plateauing to a steady state (Fig 3-FS2 left panel, similar to a reusable mediator in Fig 3-FS1A). In this case, **C_1_** may be approximated as a reusable mediator and can be predicted by the canonical pairwise model (Fig 3-FS2 right panel, compare dotted and solid lines).
Case II: When ***r*_*S*_2_*C*_1__ > *r*_10_ − *r*_20_ > 0**, a steady state solution for ***C*_1_** and ***R_S_*** = ***S*_2_** / ***S*_1_** exists (Eq.S3-4). We can rewrite

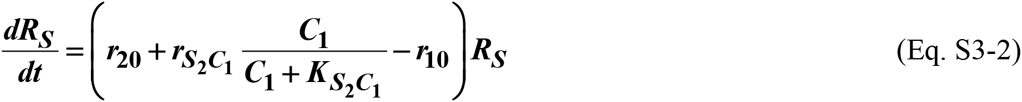

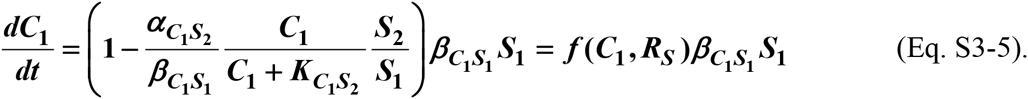 The above equations can be used to visualize the dynamics trajectory of the community in a phase plane (***C*_1_**, ***R_S_***) (Fig3-FS4, A-D). Qualitatively, starting from (**0**, ***R_S_***(**t=0**)), the trajectory moves toward the curve

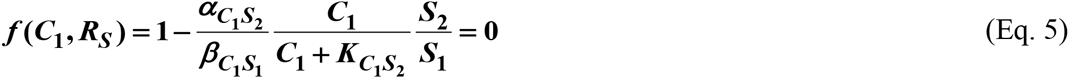

(the “***f***-zero-isocline”, blue line in Fig 3-FS4, A-D) with time scale ***t_f_*** (Fig3-FS8B third column for Case II). Afterwards the trajectory moves closely along the ***f***-zero-isocline until it reaches the steady state. During this time, we can assume ***f* ≈ 0** to eliminate ***C*_1_** and derive a pairwise model. The resultant equation below differs from the canonical pairwise model and takes the form

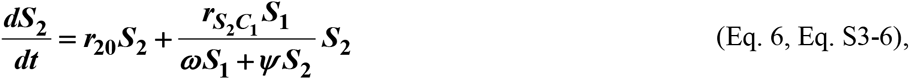

where ***ω*** and ***ψ*** are constants (Fig 3B-ii). As expected, parameter estimation for the alternative pairwise model is more accurate after the initial period of time (Fig 3-FS6 bottom panels).
Case III: When supplier **S_1_** always grows slower than consumer **S_2_** (***r*_10_ < *r*_20_**), Eqs. S3-2 and S3-5 above are still valid. Thus, consumable ***C*_1_** declines to zero concentration as it is consumed by **S_2_** whose relative abundance over **S_1_** eventually exponentially increases at a rate of ***r*_20_ − *r*_10_**. Similar to Case II, after time scale ***t_f_*** when community dynamics reaches the ***f***-zero-isocline (Fig 3-FS4 E-H; Fig3-FS8B third column for Case III), the alternative pairwise model (Eq. 6) can be derived.

The alternative pairwise model always converges to the mechanistic model if ***ω* = 1 − K_*S*_2__ / *K*_*C*_1_*S*_2__ ≥ 0** or ***K*_*C*_1_*S*_2__ ≥ *K*_*S*_2_*C*_1__** (Methods-Section 3, “Conditions for the alternative pairwise model to approximate the mechanistic model”; Fig 3-FS5A and C). Indeed, even if the initial species composition is not at steady state, the alternative model approaches the mechanistic model over time (compare dashed and solid lines in Fig 3-FS3 where ***ω* = 0**). When ***ω* < 0**, ***R_S_*(*t*=0)> −*ω*/*ψ*** is required for the alternative pairwise model to work (Fig3-FS8A). Otherwise, we can get qualitatively wrong results (Fig 3-FS5B and D).

If any of the requirements on initial conditions or time scale (Methods-Section 3; Fig3-FigS8) is violated, the alternative pairwise model may fail to represent the mechanistic model. For example, if dilution cycles are such that the community can never approach the ***f***-zero-isocline, then ***C*_1_** could accumulate proportionally to ***S*_1_** even if ***r*_*S*_2_*C*_1__ > *r*_10_ − *r*_20_ > 0**. In this case, the canonical but not the alternative pairwise model is appropriate (Fig 3-FS7). Similarly, a gentler dilution scheme, which allows the community to remain near the ***f***-zero-isocline, leads to better parameter estimation for a pairwise model (Fig 3-FS6).

The alternative model can be further simplified to

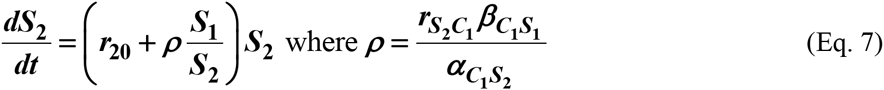

if additionally, the half-saturation constant ***K*** for **C_1_** consumption (***K*_*C*_1_*S_2_*_**) is identical to that for **C_1_**’s influence on the growth of consumer (***K*_*S*_2_*C_1_*_**) (the “***K*** assumption”), and if **S_2_** has not gone extinct. This equation form has precedence in the literature (e.g. (Mougi and Kondoh 2012)), where the interaction strength ***r*_21_** reflects the fact that the consumable mediator from **S_1_** is divided among consumer **S_2_**.

The canonical and the alternative models are not interchangeable. The alternative pairwise model is not predictive of community dynamics where **C_1_** accumulates without reaching a steady state (Fig 3-FS2, compare dashed and solid lines). Similarly, an interaction mediated by a consumable mediator that eventually reaches steady state can often be described by the alternative pairwise model (Fig 3-FS3, dashed lines). However, if we were to use a canonical model, model accuracy becomes uncertain: Parameters estimated at the steady state can predict community dynamics at steady state (Fig 3-FS3A), but not community dynamics when initial species ratios differ from the steady state ratio (Fig 3-FS3B and C, compare dotted with solid lines).

We have shown here that even when one species affects another species via a single mediator, either a canonical or an alternative form of pairwise model may be more appropriate in different situations. Choosing the appropriate model depends on whether the mediator is reusable or consumable and if consumable, how the fitness of two species compare. Additionally, a pairwise model may approximate the mechanistic model properly only under appropriate initial conditions and after the estimated time scale (Fig 3-FS5, B and D; Fig3-FS8). Considering that reusable and consumable mediators are both common, our results call for revisiting the universality assumption of pairwise modeling.

### Multi-mediator interactions require pairwise models different from single-mediator interactions

A species often affects another species via multiple mediators (Kato et al. 2008; Traxler et al. 2013; Kim, Lee, and Ryu 2013). For example, a subpopulation from one species might die and release numerous chemicals that can affect another species in different ways. Here we examine cases where **S_1_** releases two chemicals **C_1_** and **C_2_** which additively affect the growth of **S_2_** (Fig 4). We ask when two mediators can mathematically be regarded as one mediator (to facilitate further abstraction into a pairwise model), and how multi-mediator interactions affect pairwise modeling.

When both mediators are reusable (Methods-Section 4), their combined effect 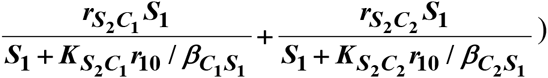 generally cannot be modeled as a single mediator except under special conditions (Fig 4). These special conditions (Methods-Section 4) include: (1) mediators share similar “potency” (Fig 4C, diagonal), or (2) one mediator has much stronger “potency” than the other (i.e. one mediator dominates the interaction; note the log scale in Fig 4C).

When both mediators are consumable as in Cases II and III, the interaction term becomes 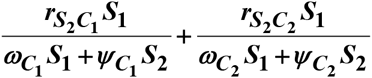. Except under special conditions (e.g. when both mediators satisfy the ***K*** assumption (Eq. 7), or when ****ω*_*C*_1__/*ω*_*C*_2__ = *Ψ*_*C*_1__/*Ψ*_*C*_2__***, or when one mediator dominates the interaction), the two mediators may not be regarded as one. Similarly, when one mediator is a steady-state consumable and the other is reusable, they generally may not be regarded as a single mediator and would require yet a different pairwise model with more degrees of freedom (with the interaction term 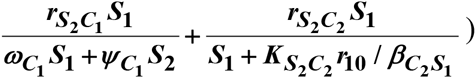). All these forms deviate from the canonical form.

In summary, when **S_1_** influences **S_2_** through multiple mediators, rarely can we approximate them as a single mediator. Multiple mediators generally make equations of pairwise modeling more complex than single mediators, casting further doubt on the usefulness of a universal form for all community interactions.

### A multispecies pairwise model can work for interaction chains but generally not for interaction modifications

For a community with more than two species, can we construct a multispecies pairwise model from two-species pairwise models? The answer is yes for an interaction chain mediated by chemicals (Fig 5A), so long as mediators between different species pairs are independent and each species pair can be represented by a pairwise model. The equation form of the multispecies pairwise model can vary (Methods-Section 5), as discussed in previous sections.

Consistent with previous work (Methods-Section 1), interaction modification can cause a multispecies pairwise model to fail. For example, **S_1_** releases **C_1_** which stimulates **S_2_** growth; **C_1_** is consumed by **S_3_** and stimulates **S_3_** growth (Fig 5C). Here, the presence of **S_3_** changes the strength of interaction between **S_1_** and **S_2_**, an example of interaction modification. Viewing this differently, **S_1_** changes the nature of interactions between **S_2_** and **S_3_**: **S_2_** and **S_3_** do not interact in the absence of **S_1_**, but **S_3_** inhibits **S_2_** in the presence of **S_1_**. This causes the three-species pairwise model to make qualitatively wrong conclusions about species persistence even though each species pair can be described by a pairwise model (Fig 5D). As expected, if **S_3_** does not remove **C_1_**, the three-species pairwise model works (Fig 5-FS1, A-B).

Interaction modification can occur even in communities where no species changes “the nature of interactions” between any other two species (Fig 5E). Here, both **S_1_** and **S_3_** contribute reusable **C_1_** to stimulate **S_2_**. **S_1_** promotes **S_2_** regardless of **S_3_**; **S_3_** promotes **S_2_** regardless of **S_1_**; **S_1_** and **S_3_** do not interact regardless of **S_2_**. However, a multispecies pairwise model assumes that the fitness effects from the two producers on **S_2_** will be additive, whereas in reality, the fitness effect on **S_2_** saturates at high ***C***_1_. As a result, even though the dynamics of each species pair can be represented by a pairwise model (Fig 5F right, purple), the three-species pairwise model fails to capture community dynamics (Fig 5F). Thus, the nonlinearity in how a mediator affects a species can also violate the additivity assumption of a pairwise model. As expected, if **C_1_** affects **S_2_** in a linear fashion, the community dynamics is accurately captured in the multispecies pairwise model (Fig 5-FS1, C-D).

In summary, for chemical-mediated indirect interactions, a multispecies pairwise model can work for the interaction chain category but generally not the interaction modification category.

## Discussion

Multispecies pairwise models are widely used in theoretical research due to their simplicity. In two-species interactions such as prey-predation based on contact-dependent inhibition (instead of diffusible chemical mediators), Lotka-Volterra pairwise models can in fact be the mechanistic representation of interactions and thus predictive of community dynamics (Fig 1-FS1). The inadequacy of multispecies pairwise models has been discussed theoretically (Wootton 2002; Wootton and Emmerson 2005) and empirically (Case and Bender 1981; Dormann and Roxburgh 2005; Aschehoug and Callaway 2015), although the reasons for model failures in explaining particular experimental results are often unclear (Billick and Case 1994).

Here, we have considered the validity of pairwise models in well-mixed two- and three-species communities where all species interactions are known and thus community dynamics can be described by a mechanistic reference model. We have focused on chemical-mediated interactions commonly encountered in microbial communities (Fig 1-FS2) (Kato et al. 2005; Gause 1934a; Ghuysen 1991; Nicholas S Jakubovics et al. 2008; Chen et al. 2004; D’Onofrio et al. 2010; Johnson et al. 1982; Hamilton and Ng 1983). To favor the odds of successful pairwise modeling, we have also assumed that different chemical mediators exert additive fitness effects on a target species.

What are the conditions under which the influence of one species on another can be represented by a canonical two-species pairwise model (the universality assumption)? When an interaction employs a single mediator, then a canonical saturable pairwise model (Fig 1B) will work for a reusable mediator (Fig 3A). For a consumable mediator, depending on the fitness relationship between the two species, either a canonical (Fig 3A) or an alternative (Fig 3B) pairwise model is appropriate (Methods-Section 3, Fig3-FS8). Canonical and alternative models are not interchangeable (Fig 3-FS2; Fig 3-FS3; Fig 3-FS7). For the alternative pairwise model, depending on the relative strength of ***K***_***C***_1_***S***_1__ and ***K***_***S***_2_***C***_1__, additional requirement on initial ***S_2_***/***S_1_*** may need to be satisfied for the pairwise model to converge to mechanistic model (Fig3-FS8A). Otherwise, qualitatively wrong predictions can ensue (Fig 3-FS5, B and D). In all cases, acclimation time is required for a pairwise model to converge to the mechanistic model (Fig3-FS8B; Fig3A). If one species influences another through multiple mediators, then in general, these mediators may not be regarded as a single mediator (Fig 4 and Methods-Section 4). Thus, conditions for a working canonical pairwise model become even more restrictive.

In communities of more than two species, indirect interactions via a third species can occur. For indirect interactions in the form of interaction chains, as long as each two-species segment of the chain engages in independent interactions and can be represented by a pairwise model, then multispecies pairwise models will generally work (Figs 5A-B, Methods-Section 5). However, depending on whether each mediator is reusable or not, equation forms will vary. For indirect interactions in the form of interaction modification (higher-order interactions), even if each species pair can be accurately represented by a pairwise model, a multispecies pairwise model may fail (Fig 5, C-F). Interaction modification includes trait modification (Wootton 2002; Werner and Peacor 2003; Schmitz, Krivan, and Ovadia 2004), or, in our cases, mediator modification. Mediator modification is very common in microbial communities. For example, antibiotic released by one species to inhibit another species may be inactivated by a third species, and this type of indirect interactions can stabilize microbial communities (Kelsic et al. 2015; Bairey, Kelsic, and Kishony 2016). Moreover, interaction mediators are often shared among multiple species. For example in oral biofilms, organic acids such as lactic acid are generated from carbohydrate fermentation by many species (Bradshaw et al. 1994; Marsh and Bradshaw 1997; Kuramitsu et al. 2007). Such by-products are also consumed by multiple species (Kolenbrander 2000).

Pairwise modeling (or variations of it) still has its uses when simulating a particular community phenomenologically. One can even imagine that an extended pairwise model (e.g. 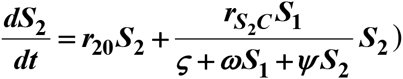) embodying both the canonical form and the alternative form can serve as a general-purpose model for pairwise interactions via a single mediator. Even the effects of indirect interactions may be quantified and included in the model by incorporating higher-order interaction terms (Case and Bender 1981; Worthen and Moore 1991), although many challenges will need to be overcome (Wootton 2002). In the end, although these strategies may lead to a sufficiently accurate phenomenological model for specific cases, “one-form-fitting-all” may generate erroneous predictions when modeling different communities.

How much information about interaction mechanisms do we need to construct a mechanistic model? That is, what is the proper level of abstraction which captures the phenomena of interest, yet avoids unnecessary details (Levins 1966; Durrett and Levin 1994; Damore and Gore 2012)? Tilman argued that if a small number of mechanisms (e.g. the “axes of trade-offs” in species’ physiological, morphological, or behavioral traits) could explain much of the observed pattern (e.g. species coexistence), then this abstraction would be highly revealing (Tilman 1987). However, the choice of abstraction is often not obvious. Consider for example a commensal community where **S_1_** grows exponentially (not explicitly depicted in equations in Fig 6) and the growth rate of **S_2_**, which is normally zero, is promoted by mediator C from **S_1_** in a linear fashion (Fig 6). If we do not know how **S_1_** stimulates **S_2_**, we can still construct a pairwise model (Fig 6A). If we know the identity of mediator **C** and realize that **C** is consumable, then we can instead construct a mechanistic model incorporating *C* (Fig 6B). However, if **C** is produced from a precursor via an enzyme **E** released by **S_1_**, then we get a different form of mechanistic model (Fig 6C). If, on the other hand, **E** is anchored on the membrane of **S_1_** and each cell expresses a similar amount of **E**, then equations in Fig 6D are mathematically equivalent to Fig 6B. This simple example, inspired by extracellular breakdown of cellulose into a consumable sugar C (Bayer and Lamed 1986; Felix and Ljungdahl 1993; Schwarz 2001)), illustrates how knowledge of mechanisms may eventually help us determine the right level of abstraction.

In summary, under certain circumstances, we may already know that interaction mechanisms fall within the domain of validity for a pairwise model. In these cases, pairwise modeling provides the appropriate level of abstraction, and constructing a pairwise model can be far easier than measuring the many parameters required by a mechanistic model. However, if we do not know whether pairwise modeling is valid, we will need to be cautious about indiscriminative use of pairwise models since they can fail to even qualitatively capture community dynamics (e.g. in Fig 3-FS2, Fig 3-FS3, Fig 3-FS5, and Fig 5C-F). We will need to be equally careful in extrapolating and generalizing conclusions obtained from pairwise models. Considering recent advances in identifying and quantifying interactions, we advocate a transition to models that incorporate interaction mechanisms at the appropriate level of abstraction.

## Methods

### Section 1. Interaction modification but not interaction chain violates the pairwise additivity assumption

In a pairwise model, the fitness of a focal species **S_i_** is the sum of its “basal fitness” (***r*_i0_**, the net growth rate of a single individual in the absence of any intra-species or inter-species interactions) and the additive fitness effects exerted by pairwise interactions with other members of the community. Mathematically, an *N*-species pairwise model is often formulated as

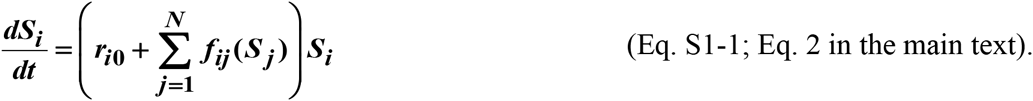

Here, *f_ij_*(*S_j_*) describes how ***S_j_***, the density of species **S_j_**, positively or negatively affects the fitness of Si, and is a linear or nonlinear function of only ***S_j_*** and not of a third species.

Indirect interactions via a third species fall under two categories (Wootton 1993). The first type is known as “interaction chain” or “density-mediated indirect interactions”. For example, the consumption of plant S1 by herbivore S2 is reduced when the density of herbivore is reduced by carnivore S3. In this case, the three-species pairwise model

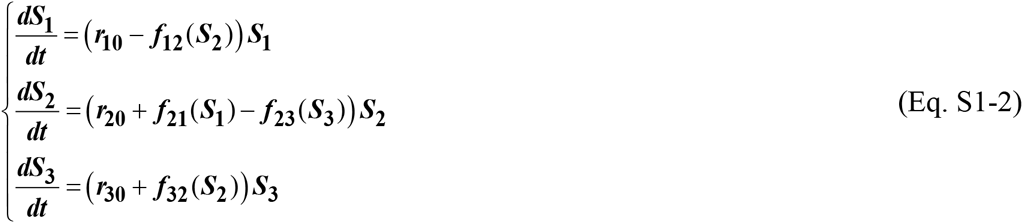

does not violate the pairwise additivity assumption (compare with Eq. S1-1) (Case and Bender 1981; Wootton 1994).

The second type of indirect interactions is known as “interaction modification” or “trait-mediated indirect interactions” or “higher order interactions” (Vandermeer 1969; Wootton 1994; Billick and Case 1994; Wootton 2002), where a third species modifies the “nature of interaction” from one species to another (Wootton 2002; Werner and Peacor 2003; Schmitz, Krivan, and Ovadia 2004). For example, when carnivore is present, herbivore will spend less time foraging and consequently plant density increases. In this case, ***f***_12_ in Eq S1-2 is a function of both ***S_2_*** and ***S_3_***, violating the pairwise additivity assumption.

### Section 2. Summary of simulation files

Simulations are based on Matlab^®^ and executed on an ordinary PC. Steps are:

Step 1: Identify monoculture parameters ***r_0_***, ***r_ii_***, and ***K_ii_*** (Fig 2C, Row 1 and Row 2).
Step 2: Identify interaction parameters ***r_ij_***, ***r_ji_***, ***K_ij_***, and ***K_ji_*** where ***i≠j*** (Fig 2C, Row 3).
Step 3: Calculate distance 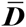 between population dynamics of the reference mechanistic model and the approximate pairwise model over several generations outside of the training window to assess if the pairwise model is predictive.

Fitting is performed using nonlinear least square (lsqnonlin routine) with default optimization parameters. The following list describes the m-files used for different steps of the analysis:

| File name | Function |
| --- | --- |
| FitCost_BasalFitness.m<br>Source code 1 | Calculates the cost function for monocultures (i.e. the difference between the target mechanistic model dynamics and the dynamics obtained from the pairwise model) |
| FitCost_BFSatLV.m<br>Source code 2 | Calculates the cost function for communities (i.e. the difference between the target mechanistic model dynamics and the dynamics obtained from the canonical pairwise model) |
| FitCost_BFSatLV_Dp.m<br>Source code 3 | Calculates the cost function for communities (i.e. the difference between the target mechanistic model dynamics and the dynamics obtained from the alternative pairwise model) |
| DynamicsMM_WM_MonocultureDpMM.m<br>Source code 4 | Returns growth dynamics for monocultures, based on the mechanistic model |
| DynamicsMMSS_WM_NetworkDpMM.m<br>Source code 5 | Returns growth dynamics for communities of multiple species, based on the mechanistic model |
| DynamicsWM_NetworkBFSatLV.m<br>Source code 6 | Returns growth dynamics for communities of multiple species, based on the canonical pairwise model |
| DynamicsWM_NetworkBFSatLV_Dp.m<br>Source code 7 | Returns growth dynamics for communities of multiple species, based on the alternative pairwise model |
| DeriveBasalFitnessMM_WM_DpMM.m<br>Source code 8 | Estimates monoculture parameters of pairwise model (Step 1) |
| DeriveBFSatLVMMSS_WM_DpMM.m<br>Source code 9 | Estimates canonical pairwise model interaction parameters (Step 2) |
| DeriveBFSatLVMMSS_Dp_WM_DpMM.m<br>Source code 10 | Estimates alternative pairwise model interaction parameters (Step 2) |
| DeriveBFSatLVMMSS_WM_DpMM_r21.m<br>Source code 11 | Estimates canonical pairwise model interaction parameters ($r_{21}$ and $K_{21}$) in cases where we know that $S_2$ is only affected by $S_1$, to accelerate optimization |
| DeriveBFSatLVMMSS_Dp_WM_DpMM_r21.m<br>Source code 12 | Estimates alternative pairwise model interaction parameter ($r_{21}$) in cases where we know that $S_2$ is only affected by $S_1$ and that $K_{S_2C_1}=K_{C_1S_2}$ to accelerate optimization |

### Section 3. Deriving a pairwise model for interactions mediated by a single consumable mediator

To facilitate mathematical analysis, we assume that requirements calculated below are eventually satisfied within each dilution cycle (see Fig3-FS7 for an example where dilution cycles violate requirements for deriving a pairwise model). We further assume *r*_10_ > 0 and *r*_20_ > 0 so that species cannot go extinct in the absence of dilution. See Fig3-FS8 for a summary of this section.

When **S_1_** releases a consumable mediator which stimulates the growth of **S_2_**, the mechanistic model as per Fig 3B, is

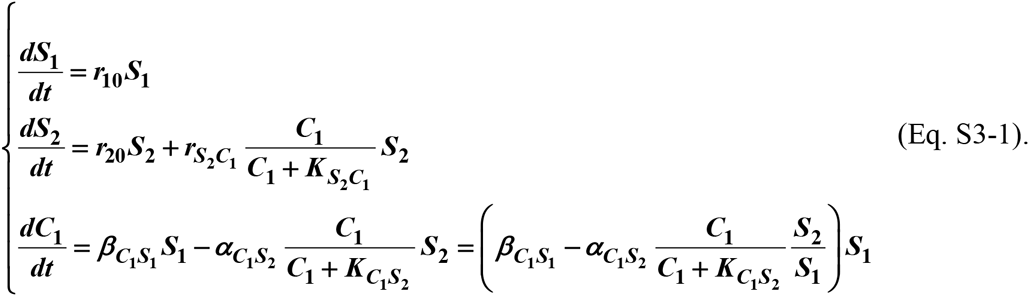

Let ***S*_1_(*t* = 0) = *S*_10_; *S*_2_(*t* = 0) = *S*_20_; *C*_1_(*t* = 0) = *C*_10_ = 0**. Note that the initial condition ***C*_10_ = 0** can be easily imposed experimentally by prewashing cells. Under which conditions can we eliminate ***C*_1_** so that we can obtain a pairwise model of ***S*_1_** and ***S*_2_**?

Define ***R_S_* = *S*_2_/*S*_1_** as the ratio of the two populations.

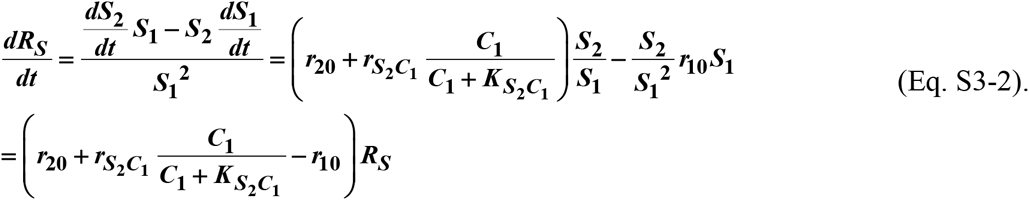

#### Case I: **r_10_ – *r*_20_ > *r*_*S*_2_*C*_1__**

Since producer **S_1_** always grows faster than consumer **S_2_**, ***R_S_*** → 0 as ***t* → ∞**. Define 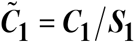 (“~” indicating scaling against a function).

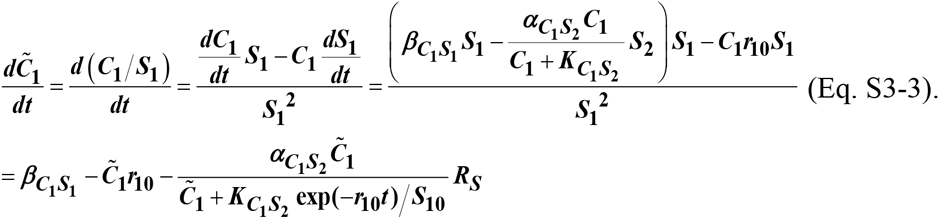

Since ***R_S_*** declines exponentially with a rate faster than 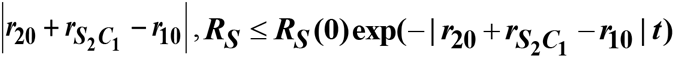. In the right hand side of Eq. S3-3, we can ignore the third term if it is much smaller than the first term. That is,

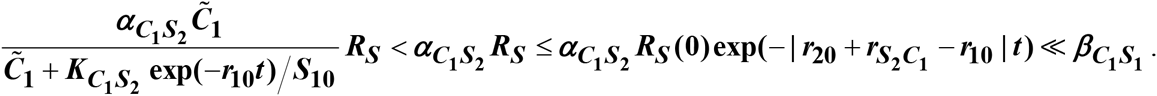

Thus for 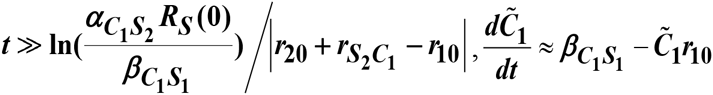. When initial 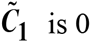, this equation can be solved to yield: 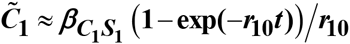. After time of the order of **1/*r_10_***, the second term can be neglected. Thus, 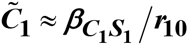 after time of the order of 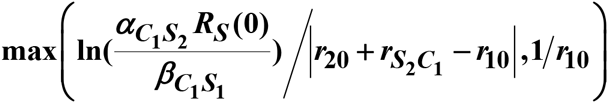. Then ***C*_1_** can be replaced by (***β*_*C*_1_*S*_1__/*r*_10_**)***S*_1_**, and a canonical pairwise model can be derived.

#### Case II: ***r*_*S*_2_*C*_1__ > *r*_10_ − *r*_20_ > 0**

For Eq. S3-1, we find that a steady state solution for ***C*_1_** and ***R_S_***, denoted respectively as ***C*_1_** * and ***R_S_***\*, exist. They can be easily found by setting the growth rates of **S_1_** and **S_2_** to be equal, and ***dC*_1_/*dt*** to zero.

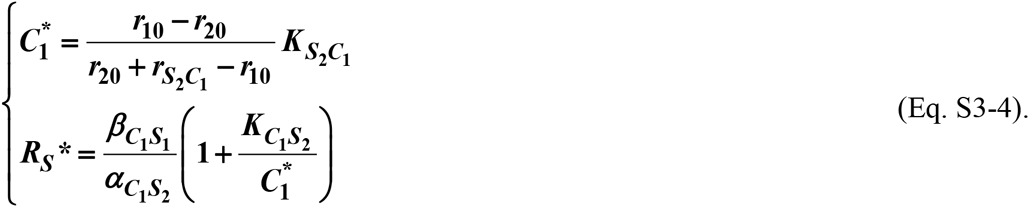

However, if ***C*_1_** has not yet reached steady state, imposing steady state assumption would falsely predict ***R_s_*** at steady state and thus remaining at its initial value (Fig 3-FS3, dotted lines). Since ***dC*_1_/*dt*** in Eq. S3-1 is the difference between two exponentially growing terms, we factor out the exponential term *S*1 to obtain

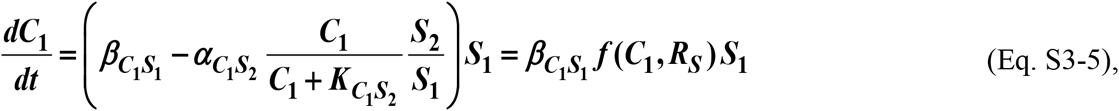

where 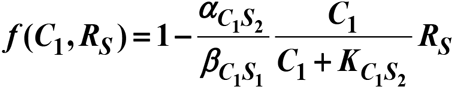. When ***f* ≈ 0**, we can eliminate ***C*_1_** and obtain an alternative pairwise model

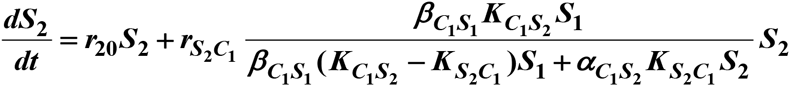

Or

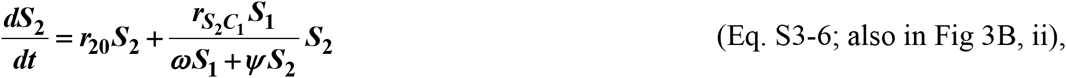

where ***ω*** and ***ψ*** are constants (Fig 3B, ii).

For certain conditions (which will be discussed at the end of this section, Fig3-FS8A), this alternative model can make reasonable predictions of community dynamics even before the community reaches the steady state (Fig 3-FS3, compare dashed and solid lines). Below we discuss the general properties of community dynamics and show that there exists a time scale *t_f_* after which it is reasonable to assume *f* ≈ 0 and the alternative model can be derived. We also estimate *t_f_* for several scenarios.

We first make ***C_1_*** and ***R_S_*** dimensionless by defining 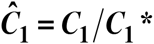 and 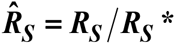 (“^” indicating scaling against steady state values). Eq. S3-2 can then be rewritten as

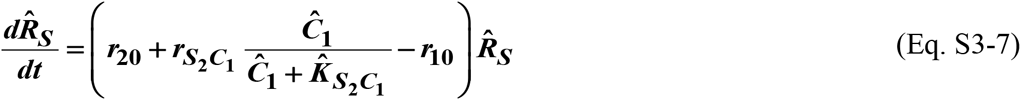

where 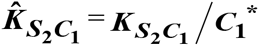.

From Eqs. S3-1 and S3-4, we obtain

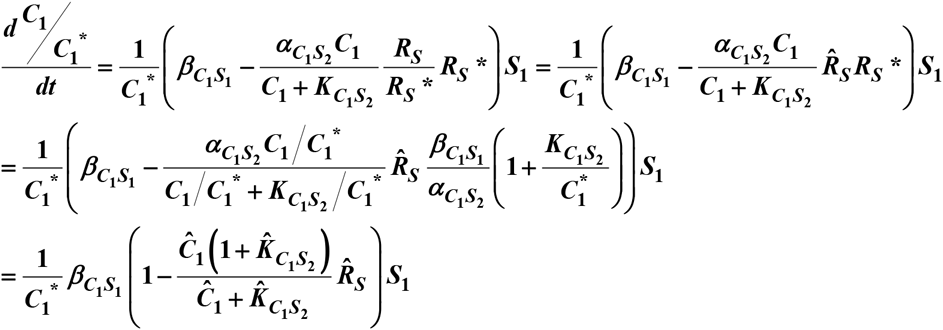

or

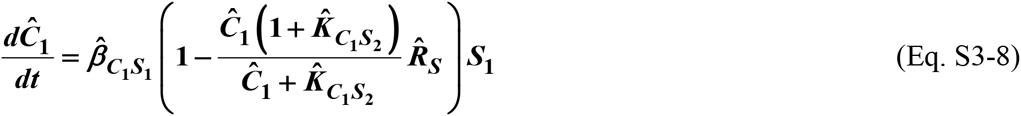

where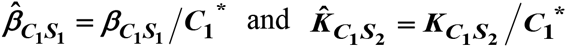.

Using these scaled variables, ***f*** can be rewritten as

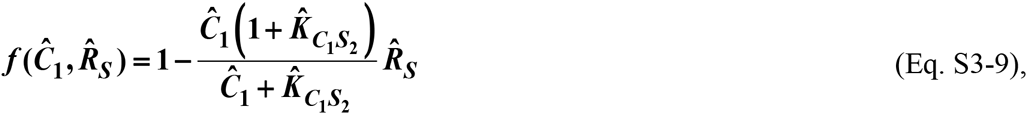

and

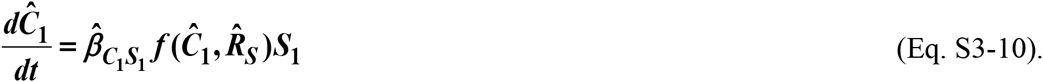

Equations S3-7 and S3-10 allow us to construct a phase portrait where the x axis is 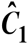 and the y axis is 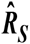 (Fig 3-FS4A-D). Note that at steady state, 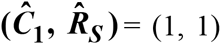. Setting Eq. S3-9 to zero:

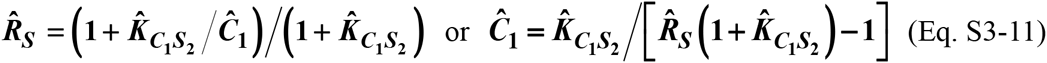

defines the ***f***-zero-isocline on the 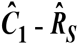 phase plane (i.e. values of 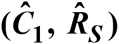 at which 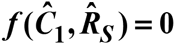, Fig 3-FS4A-D blue lines). The phase portrait dictates the direction of the community dynamics trajectory 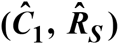 as shown in grey arrows in Fig 3-FS4A. Starting from 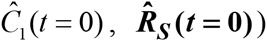, the trajectory (brown circles and orange lines) moves downward right (Fig3-FS4 A-D) until it hits 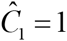. Then, it moves upward right and eventually hits the ***f***-zero-isocline. Afterward, the trajectory moves toward the steady state (green circles) very closely along (and not superimposing) the ***f***-zero-isocline during which the alternative pairwise model can be derived (Fig 3-FS4A-D).

It is difficult to solve Eq. S3-7 and S3-8 analytically because the detailed community dynamics depends on the parameters and the initial species composition in a complicated way. However, under certain initial conditions, we can estimate ***t_f_***, the time scale for the community to approach the ***f***-zero-isocline. Note that ***t_f_*** is not a precise value. Instead it estimates the acclimation time scale after which a pairwise model can be derived.

One assumption used when estimating all ***t_f_*** is that *S*_10_ is sufficiently high (Fig3-FS8B) to avoid the long lag phase that is otherwise required for the mediator to accumulate to a high enough concentration.

From Eq. S3-11, the asymptotic value for the ***t_f_***-zero-isocline is

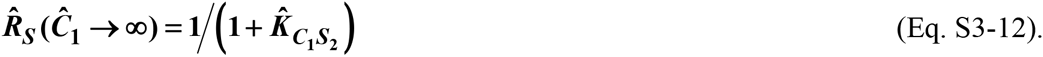

This is plotted as a black dotted line in Fig 3-FS4, A-D. Below we consider three different initial conditions 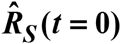:

##### Case II-1.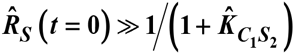

If we do not scale, 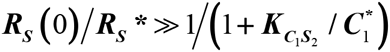. From Eq. S3-4, this becomes 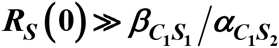.

A typical trajectory of the system is shown in Fig 3-FS4B: at time ***t = 0***, using Eqs. S3-7 and S3-8, the community dynamics trajectory (orange solid line in Fig 3-FS4B inset) has a slope of

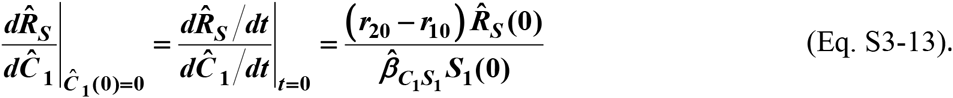

From Eqs. S3-11, the slope of the ***f***-zero-isocline (blue line in Fig 3-FS4B inset) at 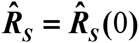 is

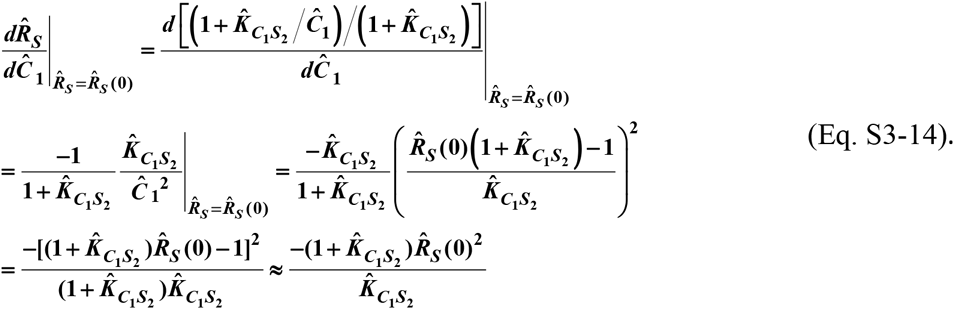

The approximation in the last step is due to the very definition of Case II-1: 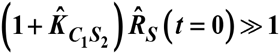. The initial steepness of the community dynamics trajectory (Eq. S3-13) will be much smaller than that of the f-zero-isocline (Eq. S3-14) if

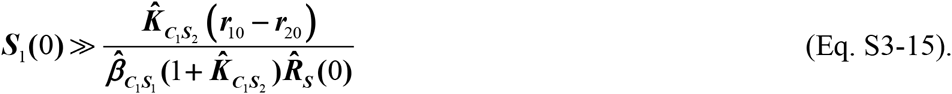

If we do not scale, together with Eq. S3-4, this becomes:

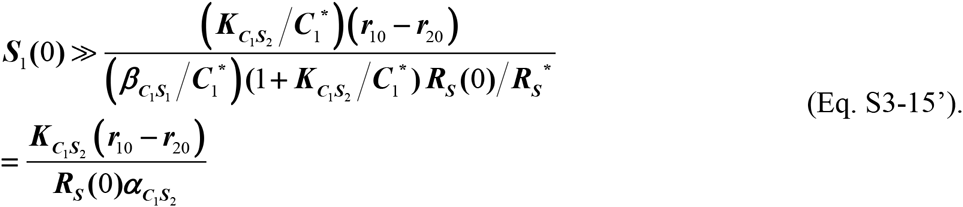

In this case, the community dynamics trajectory before getting close to the ***f***-zero-isocline can be approximated as a straight line (the orange dotted line) and the change in 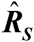 can be approximated by the green segment in the inset of Fig 3-FS4B. Since the green segment, the orange dotted line and the red dashed line form a right angle triangle, the length of green segment can be calculated once we find the length of the red dashed line 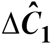, which is the horizontal distance between 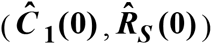 and the ***f***-zero-isocline and can be calculated from Eq. S3-9:

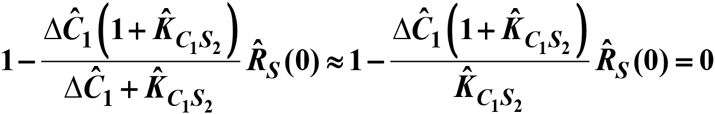

The approximation holds because under this condition, 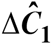 is close to zero. Thus,

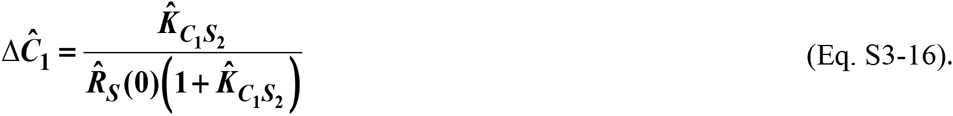

The green segment 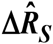 is then the length of red dashed line (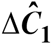, Eq. S3-16) multiplied with 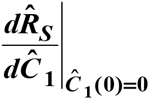 (Eq. S3-13), or

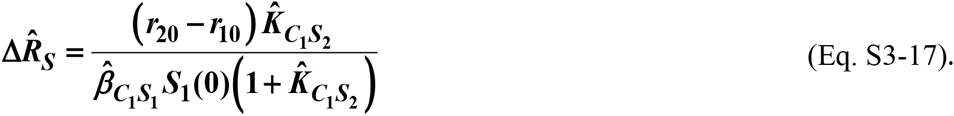

Note that if Eq. S3-15 is satisfied, 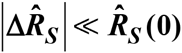. What is the time scale ***t_f_*** for the community to traverse the orange dotted line to be close to the ***f***-zero-isocline? Since from Eq. S3-7 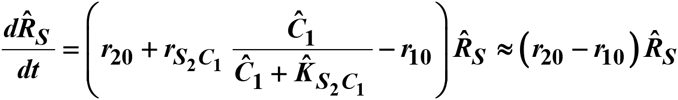 due to initial 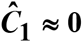,

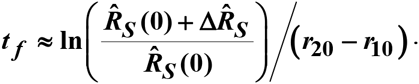

Since here 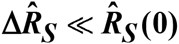 and **ln(1 + *x*) ~ *x*** for small ***x***, we have

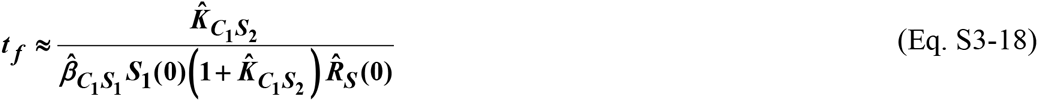

If unscaled, using Eq. S3-4, this becomes

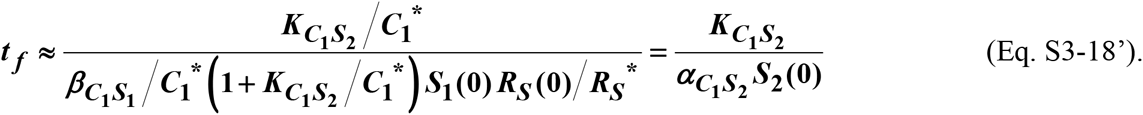

##### Case II-2. 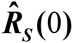 is comparable to 1

That is, 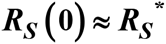. If ¾ is low, a typical example is shown in Fig 3-FS4D. Here because it takes a while for **C_1_** to accumulate, during this lagging phase 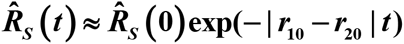 and there is a sharp plunge in 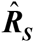 before the trajectory levels off and climbs up. Although the trajectory eventually hits the ***f***-zero-isocline where the alternative pairwise model can be derived, estimating *t_f_* is more complicated. Here we consider a simpler case where ***S*_10_** is large enough so that the trajectory levels off immediately after ***t* = 0** and 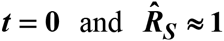 before the trajectory hits the ***f***-zero-isocline (Fig3-FS4 A). Since 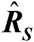 decreases until 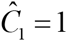 and from Eq. S3-13, and similar to the reasoning in CaseII-1, if

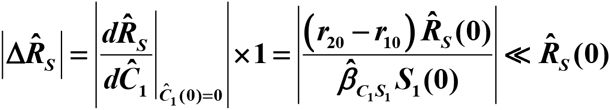

or if

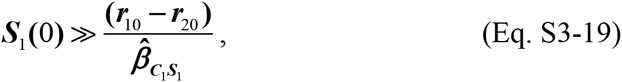

a typical trajectory moves toward the ***f***-zero-isocline almost horizontally (Fig 3-FS4A). The unscaled form of Eq. S3-19 is

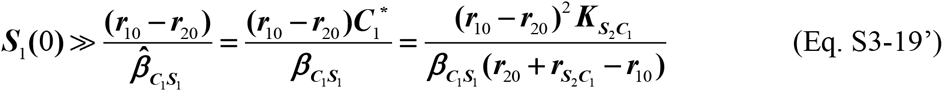

To calculate the time it takes for the trajectory to reach the ***f***-zero-isocline, let 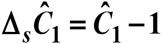 and 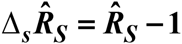 at any time point ***t*** to respectively represent deviation of 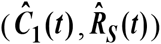 away from their steady state values of (1, 1). We can thus linearize Eqs. S3-7 and S3-8 around the steady state. Note that at the steady state ***f*** = **0**, Δ***_s_f = f***.

Rewrite Eq. S3-7 as

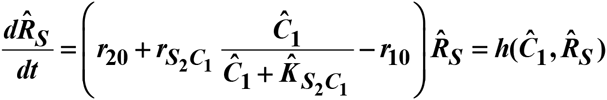

to linearize it around the steady state 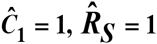

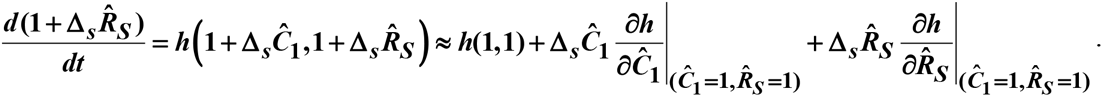

At steady state, 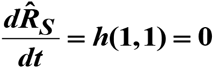. Thus, 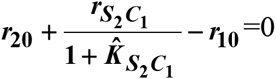.

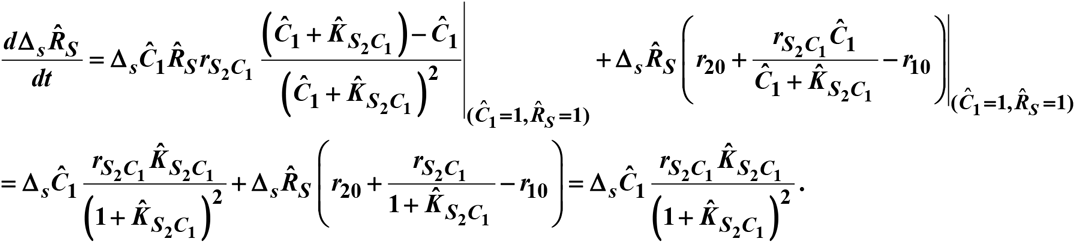

Thus,

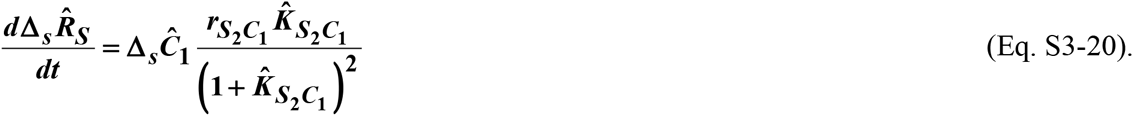

Recall Eqs. S3-8 and S3-10 as

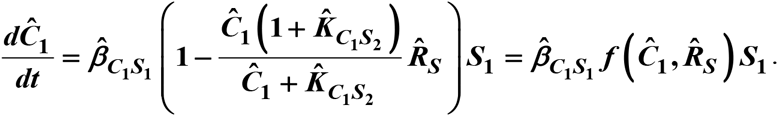

Linearize around the steady state 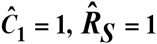 (note ***f* (1,1)=0**):

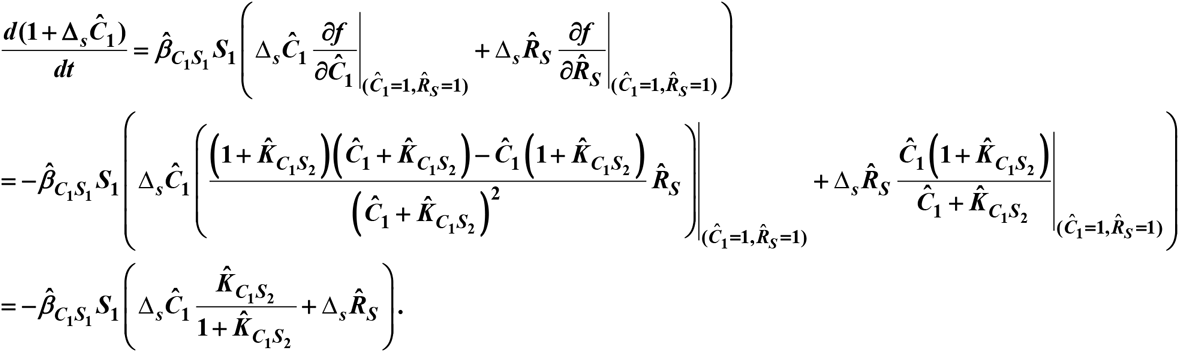

Thus,

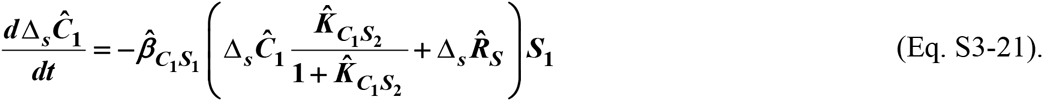

Similar to the above calculation, we expand ***f*** around steady state 0,

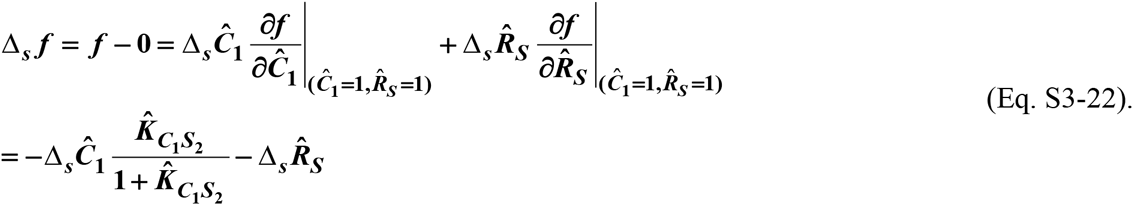

Utilizing Eq. S3-20, Eq. S3-21, and Eq. S3-22,

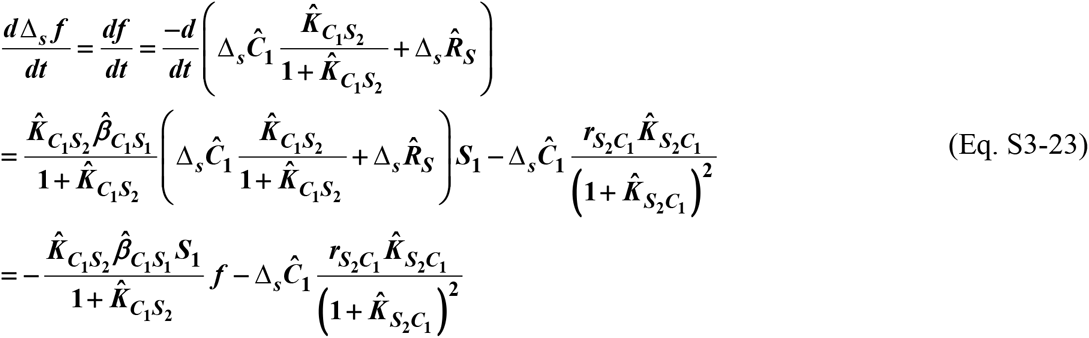

Taking the derivative of both sides, and using Eq. S3-21 and Eq. S3-22, we have

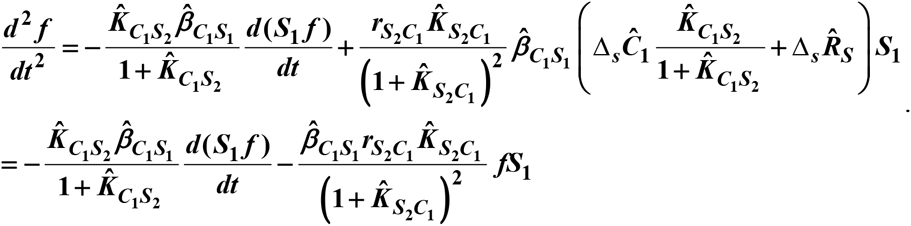

The solution to the above equation is:

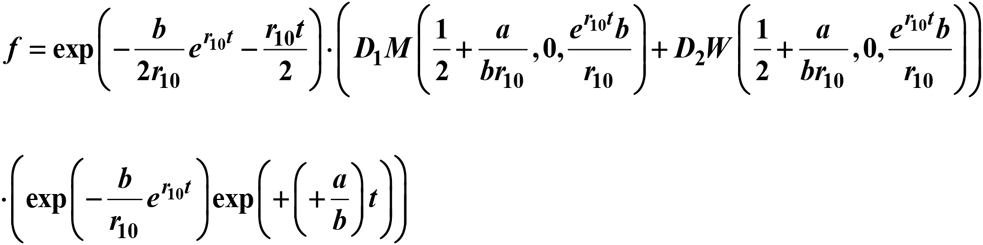

where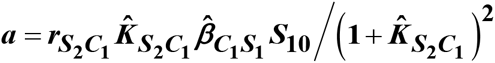 and 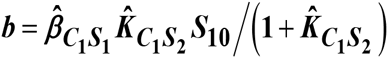 are two positive constants. ***D*_1_** and ***D*_2_** are two constants that can be determined from the initial conditions of 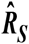 and 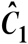. ***M*(*κ*, *μ*, *z*)** and ***W*(*κ*, *μ*, *z*)** are Whittaker functions with argument ***z***. As ***Z*** → ∞ (http://dlmf.nist.gov/13.14.E20 and http://dlmf.nist.gov/13.14.E21)

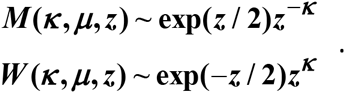

Thus when ***e*^*r*_10_*t*^*b*/*r*_10_» 1**,

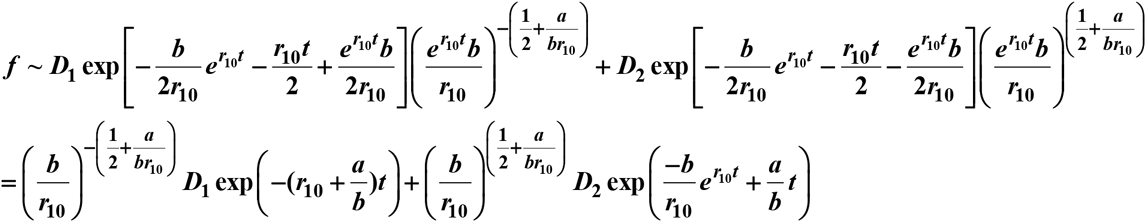

The second term approaches zero much faster compared to the first term due to the negative exponent with an exponential term. Thus,

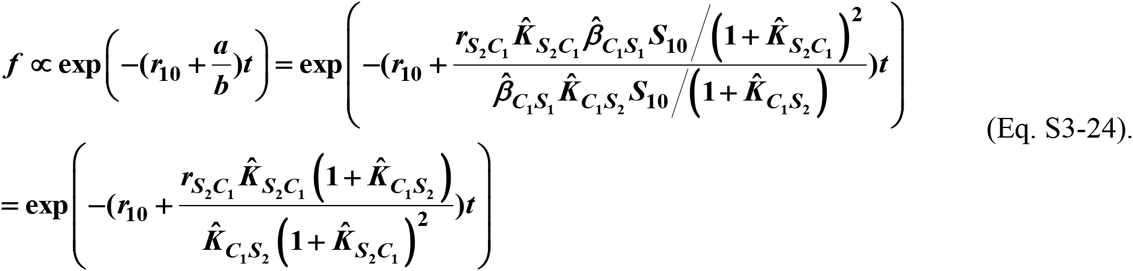

Thus, Δ***_s_f = f*** approaches zero at a rate of 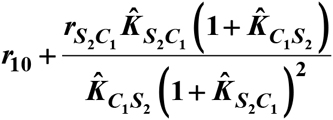. Therefore, for

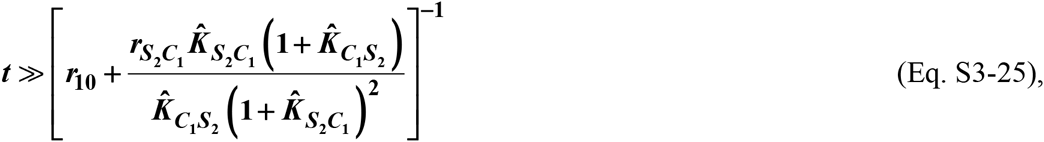

the community is sufficiently close to ***f***-zero-isocline.

In unscaled form, this becomes:

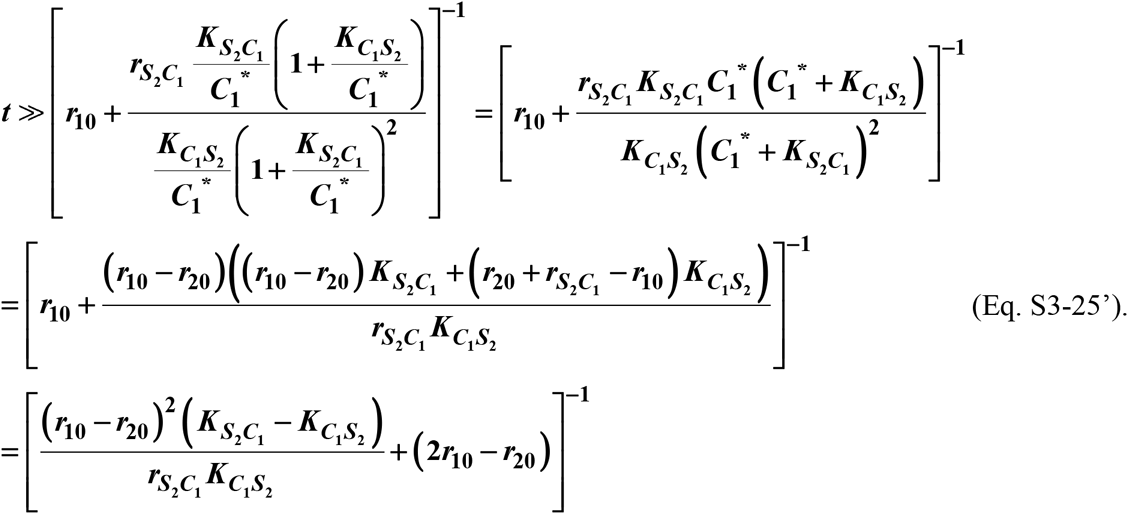

##### Case II-3.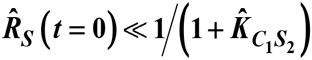 or ***R_S_* (0) ≪ *β*_*C*_1_*S*_1__/*α*_*C*_1_*S*_2__**

Similar to Case II-2, if Eq. S3-19 is satisfied, a typical trajectory is illustrated in Fig 3-FS4C where the trajectory is almost horizontal and 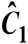 increases to much greater than 1 before the system reaches the ***f***-zero-isocline. ***t_f_*** can then be estimated from how long it takes 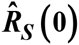 to increase to 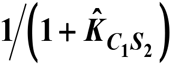. Using Eq. S3-7 and since 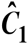 is very large,

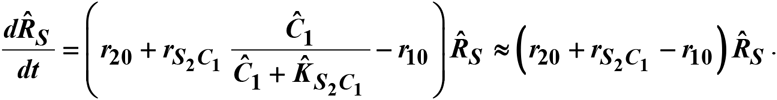

This yields

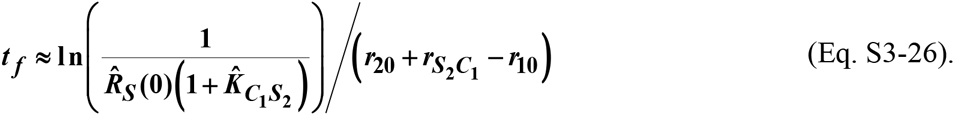

In the unscaled form, this becomes:

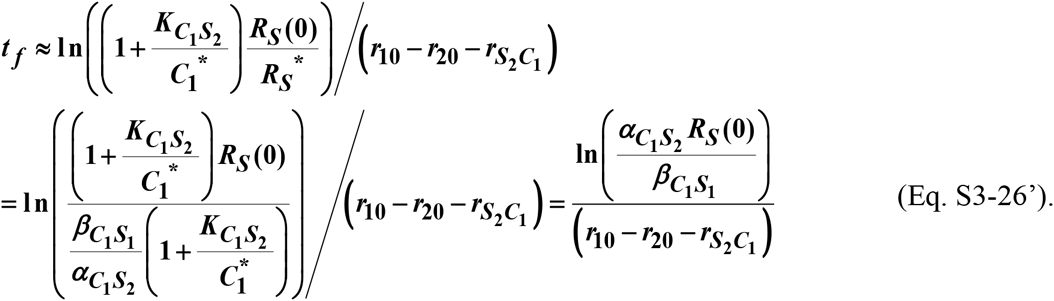

#### Case III: ***r*_20_ < *r*_20_**

In this case, supplier **S_1_** always grows slower than **S_2_**. As ***t*** → ∞, ***R_S_* = *S*_2_/*S*_1_ → ∞** and ***C*_1_ → 0**. The phase portrait is separated into two parts by the ***f***-zero-isocline (Fig 3-FS4E), where, as in Eq. S3-5,

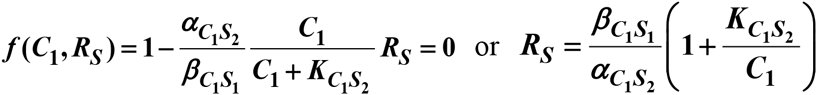

Note that the asymptotic value of ***R_S_*** (black dotted line, Fig 3-FS4, E-H) is

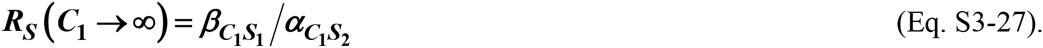

From Eq. S3-2, ***dR_S_*/*dt* > 0**. From Eq. S3-1, below the ***f***-zero-isocline, ***dC*_1_/*dt* > 0** and above the ***f***-zero-isocline, ***dC*_1_/*dt* > 0**. Thus if the system starts from (0, ***R_s_*** (0)), the phase portrait dictates that it moves with a positive slope until a time of a scale *t_f_* when it hits the ***f***-zero-isocline, after which it moves upward to the left closely along the ***f***-zero-isocline (Fig3-FS4E). After ***t*_f_**, the alternative pairwise model can be derived. Although ***t_f_*** is difficult to estimate in general, it is possible for the following cases.

##### Case III-1. ***R_S_*(0)≫*β*_*C*_1_*S*_1__/*α*_*C*_1_*S*_2__**

Similar to the derivation in Case II-2, if ***S*_10_** is small, there is a lagging phase during which the trajectory rises steeply before leveling off (Fig 3-FS4H). Although the alternative pairwise model can be derived once the trajectory hits the ***f***-zero-isocline, *t_f_* takes a complicated form. Here we consider two cases where S_10_ is large enough so that we can approximate the trajectory as a straight line going through (0, ***R_S_***(***t***=0)) (Fig3-FS4, F, G). Graphically, £_10_ is large enough so that the green segment in Fig3-FS4F, whose length is Δ***R_S_***, is much smaller than *R_S_*(0). In other words,

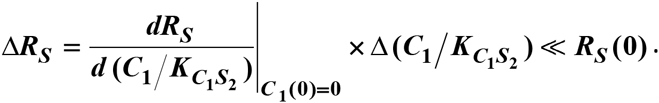

From Eqs. S3-1 and S3-2

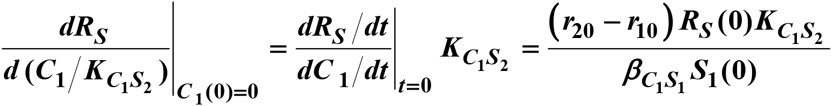

And Δ**(*C*_1_/*K*_*C*_1_*S*_2__)**, the red segment in Fig3-FS4F, is the horizontal distance between (0, ***R_S_*** (0)) and the ***f***-zero-isocline and

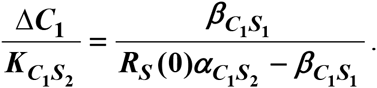

Thus if

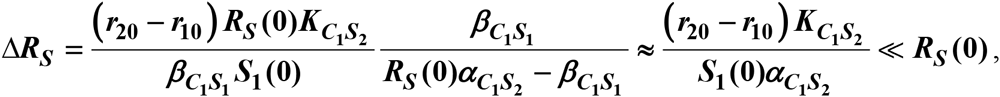

or

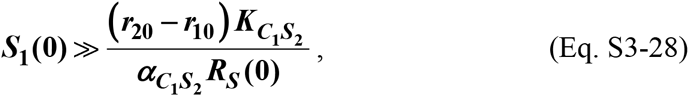

from Eq. S3-2, ***t_f_*** can be calculated as

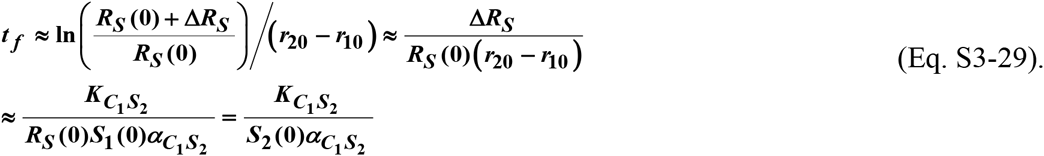

##### Case III-2. ***R_S_*(0)≫*β*_*C*_1_*S*_1__/*α*_*C*_1_*S*_2__**

If *S*_10_ is large enough so that Eq. S3-28 is satisfied, a typical example is displayed in Fig3-FS4G. The trajectory moves with a small positive slope so that the intersection of the community dynamics trajectory with the ***f***-zero-isocline is near the black dotted line ***β*_*C*_1_*S*_1__/*α*_*C*_1_*S*_2__** (Eq. S3-27) where ***C*_1_/*K*_*C*_1__** is large. ***t_y_*** can thus be estimated from Eq. S3-2:

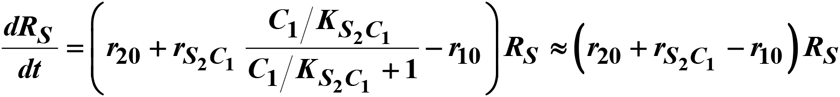

which yields

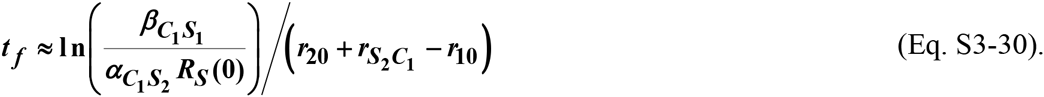

#### Conditions for the alternative pairwise model to approximate the mechanistic model

Cases II and III showed that population dynamics of the mechanistic model could be described by the alternative pairwise model. However, since the initial condition for ***C*_1_** cannot be specified in pairwise model, problems could occur. To illustrate, we examine the phase portrait of the pairwise equation

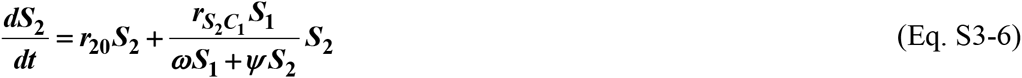

where 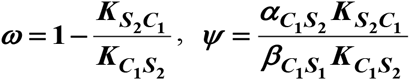. From Eqs. S3-6 and S3-1,

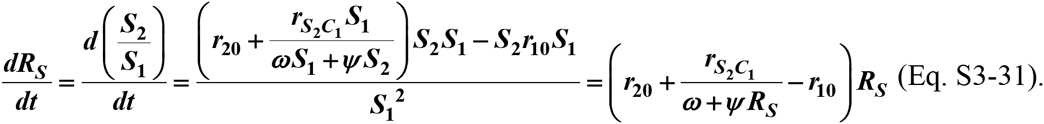

Below, we plot Eq. S3-31 under different parameters (Fig 3-FS5) to reveal conditions for convergence between mechanistic and pairwise models.

- Case II (***r*_*S*_2_*C*_1__ >*r*_10_ − *r*_20_ > 0**): steady state ***R_S_***\* exists for mechanistic model. If ***ω* = 1− *K*_*S*_2_*C*_1__/*K*_*C*_1_*S*_2__ ≥ 0**(Fig 3-FS5A): When **R_S_** <***R_S_***\*, ***dR_S_*/*dt*** is positive. When ***Rs*** > ***R_S_***\*, ***dR_S_*/*dt*** is negative. Thus, wherever the initial ***R_S_***, it will always converge toward the only steady state ***R_S_***\* of the mechanistic model. If ***ω* < 0** (Fig 3-FS5B): ***ω + ψR*_*S*_ = 0** or ***R_S_* = −*ω/ψ*** creates singularity. Pairwise model ***R_S_*** will only converge toward the mechanistic model steady state if

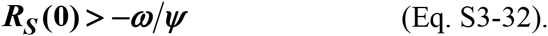
- Case III (***r*_10_ < *r*_20_**): ***R_S_*** increases exponentially in mechanistic model (Eq. S3-2). Thus, consumable ***C*_1_** will decline toward zero as **C_1_** is consumed by **S_2_** whose relative abundance over **S_1_** exponentially increases. Thus, from mechanistic model (Eq. S3-2), ***R_S_*** eventually increases exponentially at a rate of ***r*_20_ − *r*_10_**. If ***ω* ≥ 0** (Fig 3-FS5C): Eq. S3-31 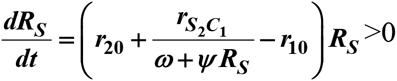. Thus, Eq. S3-31, which is based on alternative pairwise model, also predicts that ***R_S_*** will eventually increase exponentially at a rate of ***r*_20_ − *r*_10_**, similar to the mechanistic model. If ***ω* < 0** (Fig 3-FS5D): ***R_S_* (0) > −ω/ψ** (Eq. S3-32) is required for unbounded increase in ***R_S_***(similar to the mechanistic model). Otherwise, ***R_S_*** converges to an erroneous value instead.

### Section 4. Conditions under which a pairwise model can represent one species influencing another via two reusable mediators

Here, we examine a simple case where S1 releases reusable **C_1_** and **C_2_**, and **C_1_** and **C_2_** additively affect the growth of S2 (see example in Fig 4).

The mechanistic model is:

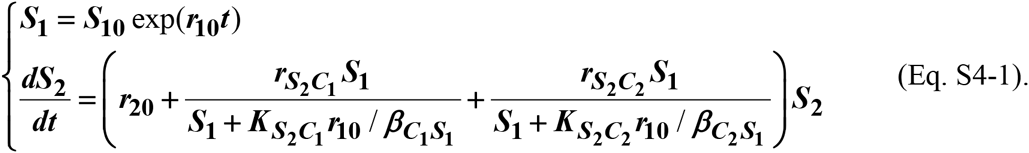

Now the question is whether the canonical pairwise model

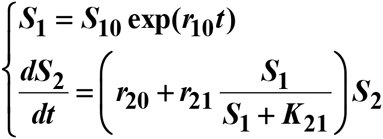

can be a good approximation.

For simplicity, let’s define 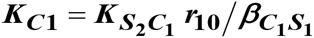 and 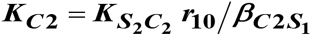. Small ***K_C_i__*** means large potency (e.g. small ***K*_*S*_2_*C*_2__** which means low ***C_2_*** required to achieve half maximal effect on ***S*_2_**, and/or large synthesis rate *β*_*C*_2_*S*_1__). Since ***S*_1_** from pairwise and mechanistic models are identical, we have

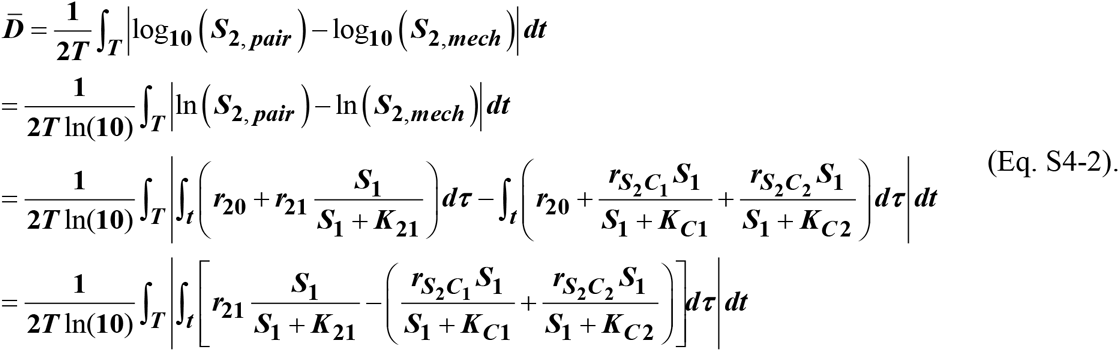

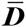 can be close to zero when (i) 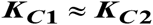 or (ii) 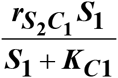 and 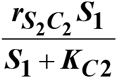 (effects of **C_1_** and **C_2_** on **S_2_**) differ dramatically in magnitude. For (ii), without loss of generality, suppose that the effect of **C_2_** on **S_2_** can be neglected. This can be achieved if (iia) ***r*_*S*_2_*C*_2__** is much smaller than ***r*_*S*_2_*C*_2__**, or (iib) ***K*_*C*_2__** is large compared to ***S_1_***.

### Section 5. Multi-species pairwise model for an interaction chain

Without loss of generality, we consider an example where each step of an interaction chain is best represented by a different form of pairwise model. Suppose that **S_1_** releases a reusable mediator **C_1_** that promotes **S_2_** and that **S_2_** releases a consumable mediator **C_2_** that promotes **S_3_**. The interaction mediated by **C_2_** can be described by the alterative pairwise model. The corresponding community-pairwise model then will be:

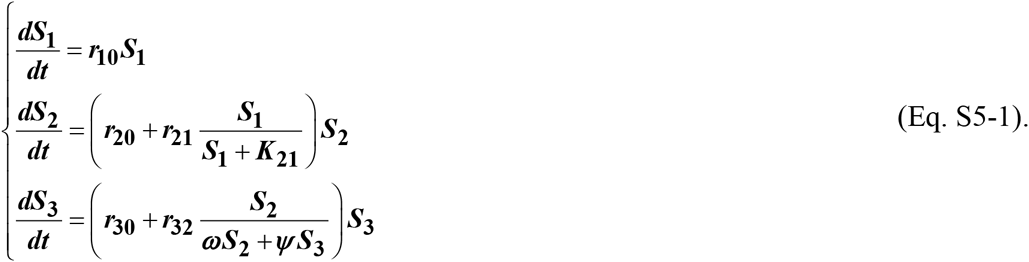

To see this, note that after a transient period of time, the mechanistic model of the three-species community is:

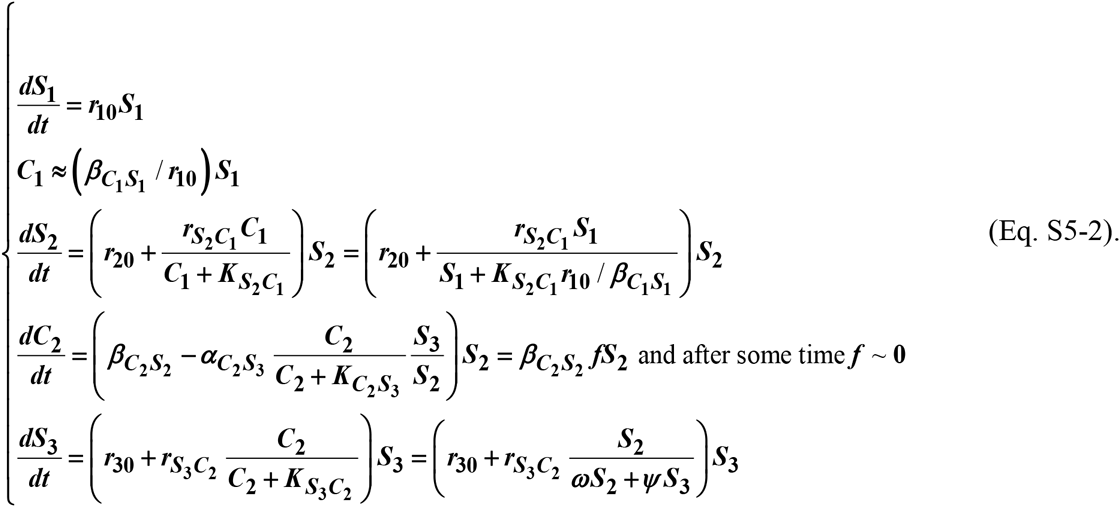

Equations S5-2 and S5-1 are equivalent.

## Figures

**Fig 1.**
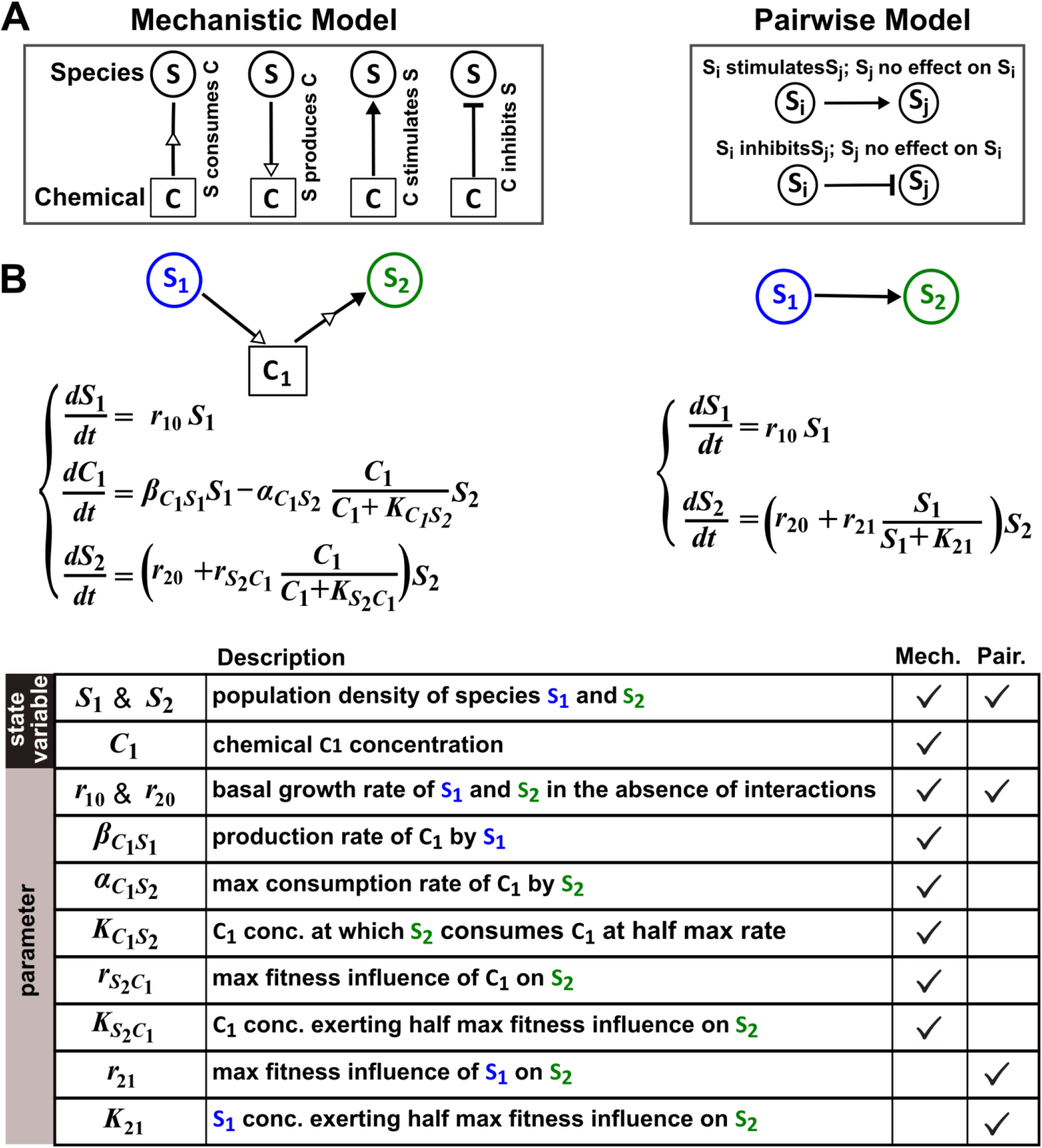
The abstraction of interaction mechanisms in a pairwise model compared to a mechanistic model. (A) The mechanistic model (left) considers a bipartite network of species and chemical interaction mediators. A species can produce or consume chemicals (open arrowheads pointing towards and away from the chemical, respectively). A chemical mediator can positively or negatively influence the fitness of its target species (filled arrowhead and bar, respectively). The corresponding pairwise model (right) includes only the fitness effects of species interactions, which can be positive (filled arrowhead), negative (bar), or zero (line terminus). (B) In the example here, species **S_1_** releases chemical **C_1_**, and **C_1_** is consumed by species **S_2_** and promotes **S_2_**’s fitness. In the mechanistic model, the three equations respectively state that 1) **S_1_** grows exponentially at a rate ***r_10_***, 2) **C_1_** is released by **S_1_** at a rate ***β*_*C*_1_*S*1_** and consumed by **S_2_** with saturable kinetics, and 3) S2’s growth (basal fitness ***r_20_***) is influenced by *Ci* in a saturable fashion. In the pairwise model here, the first equation is identical to that of the mechanistic model. The second equation is similar to the last equation of the mechanistic model except that ***r_21_*** and ***K_21_*** together reflect how the density of **S_1_** (***S_1_***) affects the fitness of **S_2_** in a saturable fashion. For all parameters with double subscripts, the first subscript denotes the focal species or chemical, and the second subscript denotes the influencer. Note that unlike in mechanistic models, we have omitted “***S***” from subscripts in pairwise models (e.g. ***r_21_*** instead of ***r*_*S*_2_*S*_1__**) for simplicity.

**Fig 2.**
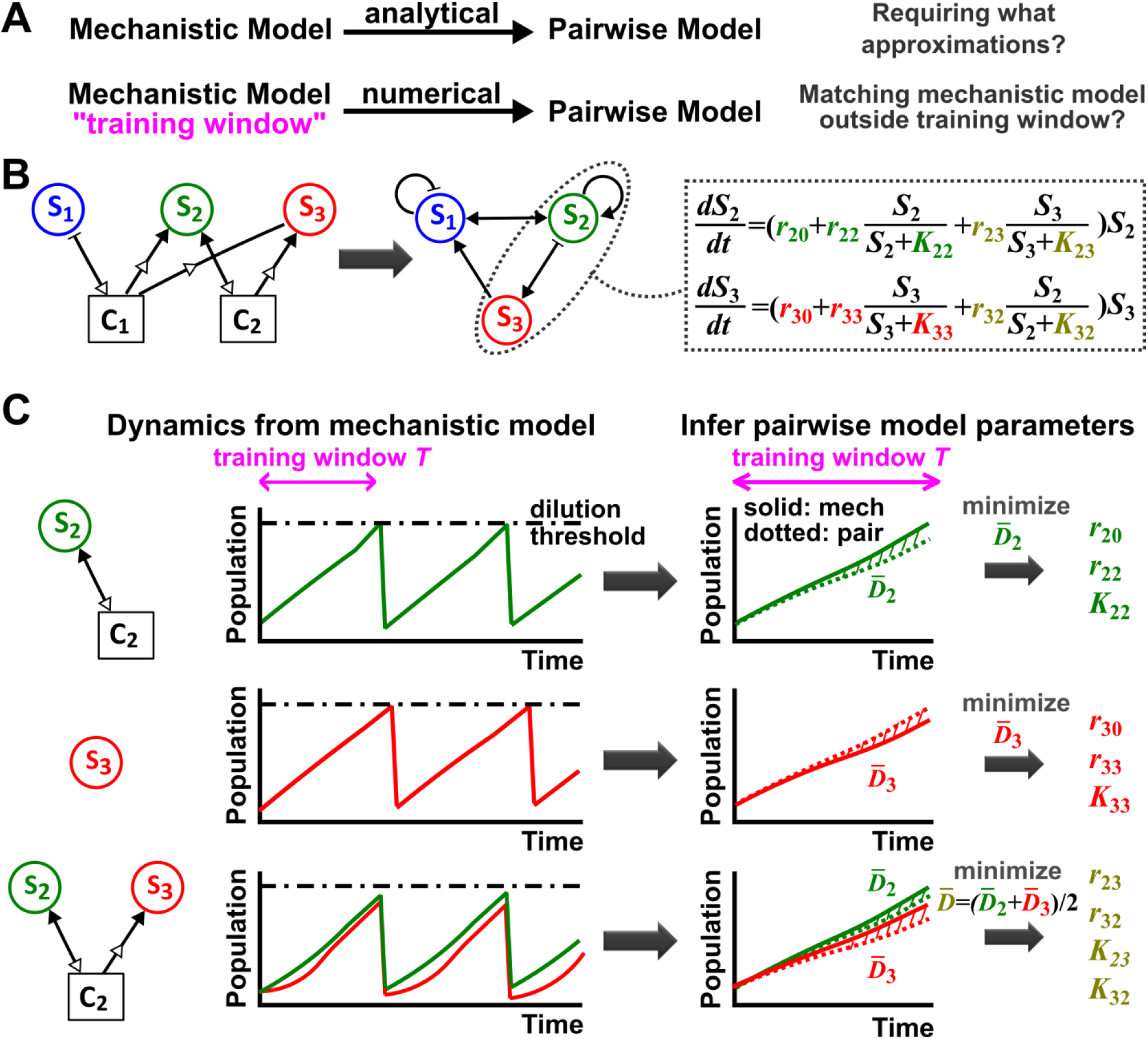
Deriving a pairwise model. (A) Analytically deriving a pairwise model from a mechanistic model allows us to uncover approximations required for such a transformation (top). Alternatively (bottom), through a “training window” of the mechanistic model population dynamics, we can numerically derive parameters for a preselected pairwise model that best fits the mechanistic model. We then quantify how well such a pairwise model matches the mechanistic model under conditions different from those of the training window. (**B**) A mechanistic model of three species interacting via two chemicals (left) can be translated into a pairwise model of three interacting species (center). **S_1_** inhibits **S_1_** and promotes **S_2_** (via **C_1_**). **S_2_** promotes **S_2_** and **S_3_** (via C2) as well as **S_1_** (via removal of **C_1_**). S3 promotes **S_1_** and inhibits **S_2_** (via removal of **C_1_** and **C_2_**, respectively). Take interactions between **S_2_** and **S_3_** for example: the saturable Lotka-Volterra pairwise model will require estimating ten parameters (colored, right), some of which (e.g. ***r_33_*** in this case) may be zero. (**C**) In the numerical method, the six monoculture parameters (***r_i0_***, ***r_ii_***, and ***K_ii_***, ***i***=2, 3; green and red) are first estimated from training window *T* (within a dilution cycle) of monoculture mechanistic models. subsequently, the four interaction parameters (***r_ij_*** and ***K_ij_, i***_≠_***j***, olive) can be estimated from the training window ***T*** of the ***S_2_*** + ***S_3_*** coculture mechanistic model. Parameter definitions are described in Fig 1. To estimate parameters, we use an optimization routine to minimize 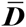, the fold-difference (shaded area) between dynamics from a pairwise model (dotted lines) and the mechanistic model (solid lines) averaged over ***T***. In all simulations, to ensure that resources not involved in interactions are never limiting, a community is diluted back to its inoculation density when total population increases to a high-density threshold. Too frequent dilutions will allow only small changes in population dynamics within a dilution cycle or ***T***, which is not suitable for estimating pairwise models. Too infrequent and large dilutions will cause large fluctuations in dynamics, which can sometimes violate conditions for deriving pairwise models (Fig 3-FS6; Fig 3-FS7). Under most cases we have tested, small variations in dilution frequency do not affect our conclusions. See Methods-Section 2 for relevant Matlab codes.

**Fig 3.**
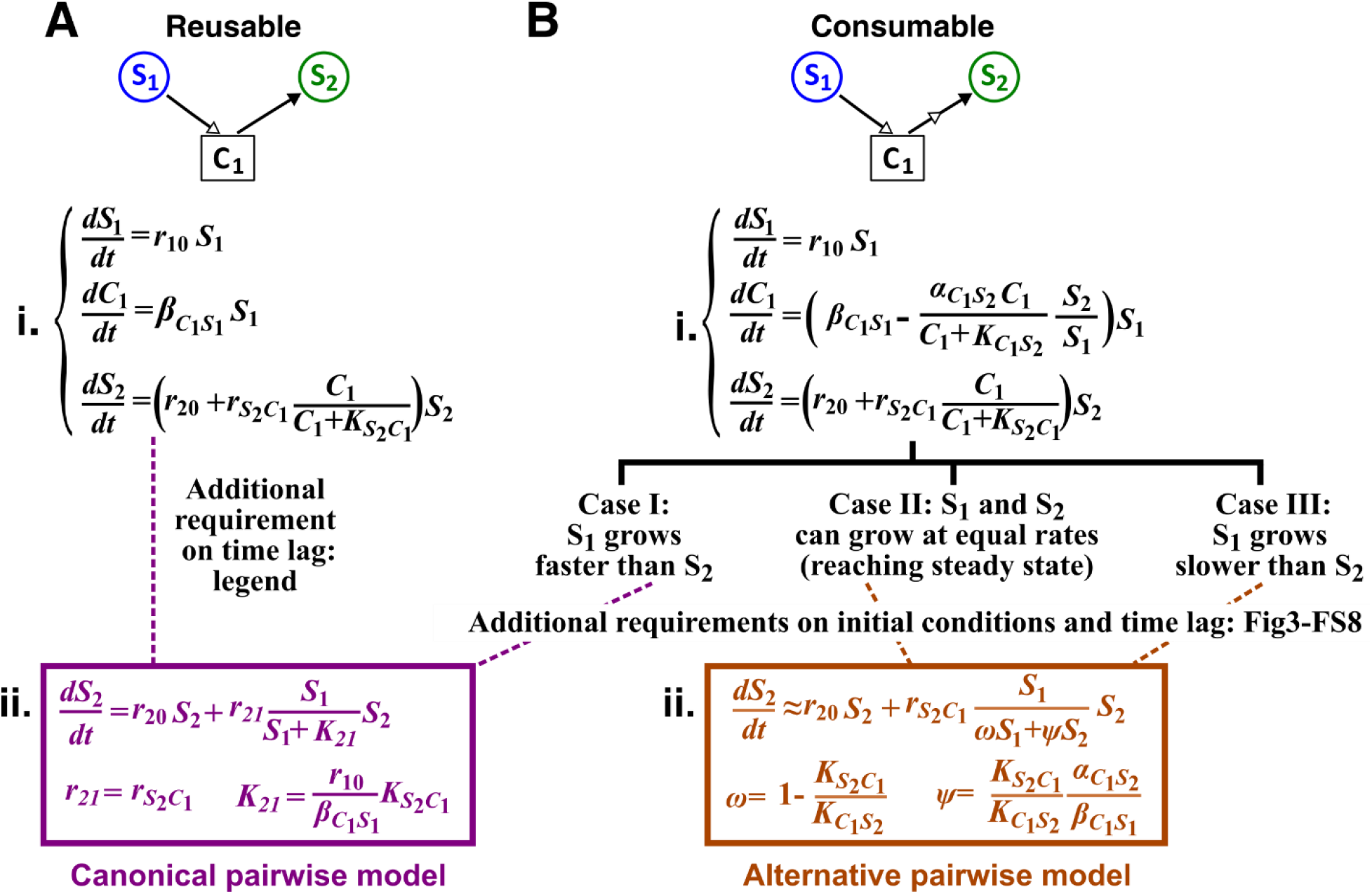
Interactions mediated via a single reusable or consumable mediator are best represented by different forms of pairwise models. **S_1_** stimulates the growth of **S_2_** via a reusable (**A**) or a consumable (**B**) chemical **C_1_**. In mechanistic models of the two cases, equations for ***S_1_*** and ***S_2_*** are identical but equations for ***C_1_*** are different. In (**A**), ***C*_1_** can be solved to yield 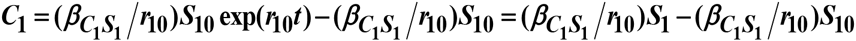 assuming zero initial ***C*_1_**. We have approximated **C_1_** by omitting the second term (valid after the initial transient response has passed so that ***C*_1_** has become proportional to ***S_1_***). This approximation allows an exact match between the canonical pairwise model (Fig 1B, right) and the mechanistic model (ii), and thus justifies the pairwise model. In (**B**), depending on the relative growth rates of the two species, and if additional requirements are satisfied (Methods-Section 3; Fig 3-FS7; Fig 3-FS8), canonical or alternative pairwise model should be used. Thus, depending on whether the mediator is consumed or reused, the most appropriate pairwise model (colored) takes different forms.

**Fig 4.**
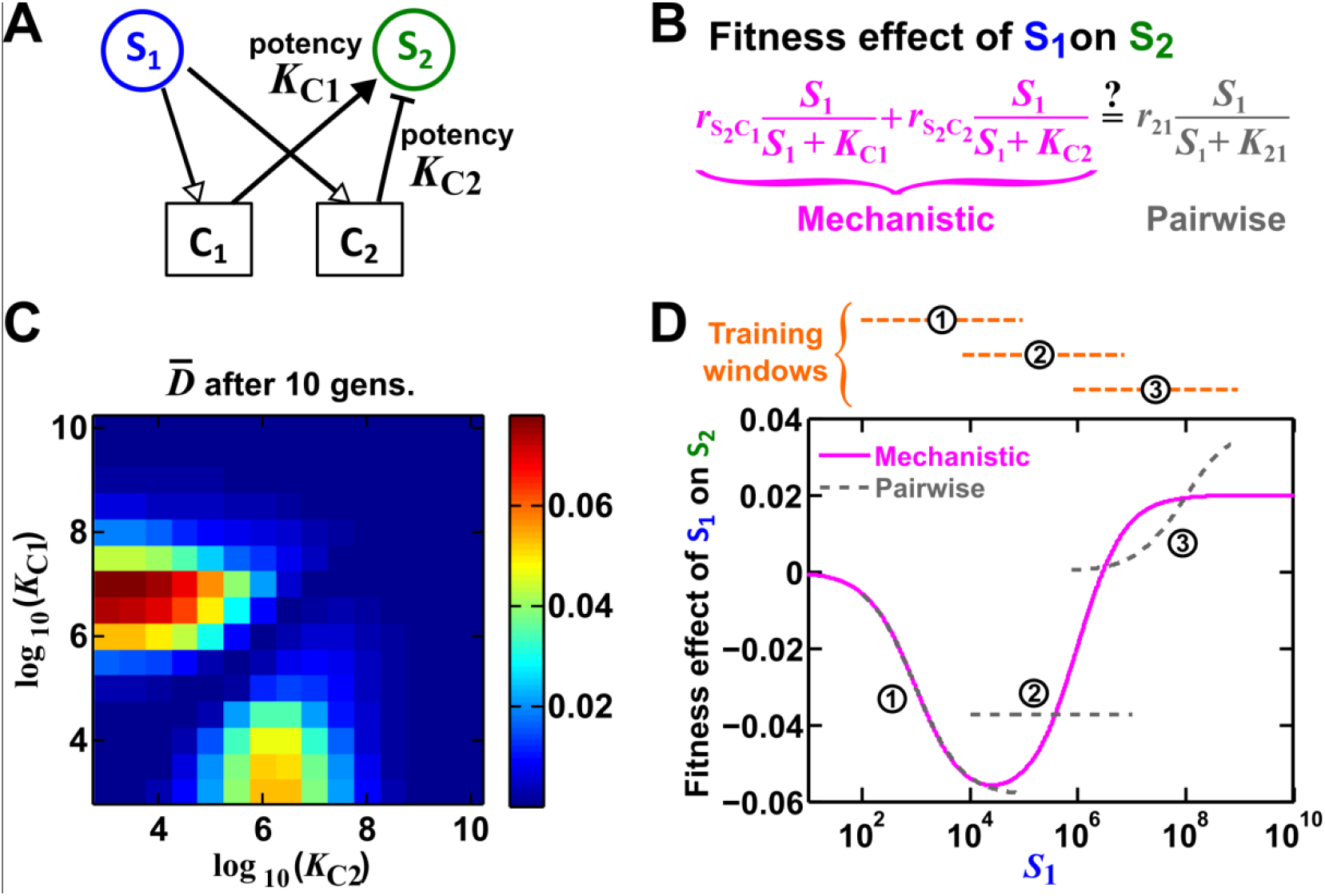
A pairwise model often fails when one species affects another via multiple reusable mediators. (**A**) One species can affect another species via two reusable mediators, each with a different potency ***K_*C*_i__*** where *Ka* is ***K*_S2_*C*_i_ *r*_10_/*β_S_i__S*_1_** (Methods-Section 4). A low ***K_C_i__*** indicates a strong potency (e.g. high release of **C_i_** by **S_1_** or low ***C_i_*** required to achieve half-maximal influence on **S_2_**). (**B**) Under what conditions can an interaction via two reusable mediators be approximated by a pairwise model? (**C**) Under restricted conditions, two reusable mediators can be consolidated into a single mediator. We can directly compute the best-fitting pairwise model parameters over a training window of ***T*** by minimizing 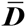 (Methods-Section 4, Eq. S4-2). Here, the difference 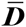 between the two models over ***T*** =**10** generations is plotted over a range of potencies ***K_C_1__*** and ***K_C_2__***. The canonical pairwise model is valid (blue regions indicating small difference) when ***K_C_1__***≈***K_C_2__*** or when one interaction is orders of magnitude stronger than the other interaction (Methods-Section 4). (**D**) A community where the canonical pairwise model is not valid. Here, ***K_C_1__***=10^3^ and ***K_C_2__***=10^6^. We estimate the best-fitting pairwise model by minimizing *D* (Methods-Section 4, Eq. S4-2) in three training windows (spanning 10 generations of growth for **S_1_**). At various ***S_1_***, we calculate the fitness effect of **S_1_** on **S_2_** using the pairwise model and the mechanistic model (B). In two of the three training windows, the two models fail to match. In the training window with the lowest *Si*, the two models match because the effect of **C_2_** is negligible in this range (***K*_*C*_2__>>*S*_1_**, condition iib in Methods-Section 4). These mismatches mean that a pairwise model cannot consistently capture reference dynamics. Simulation parameters are listed in Fig 4-SD1.

**Fig 5.**
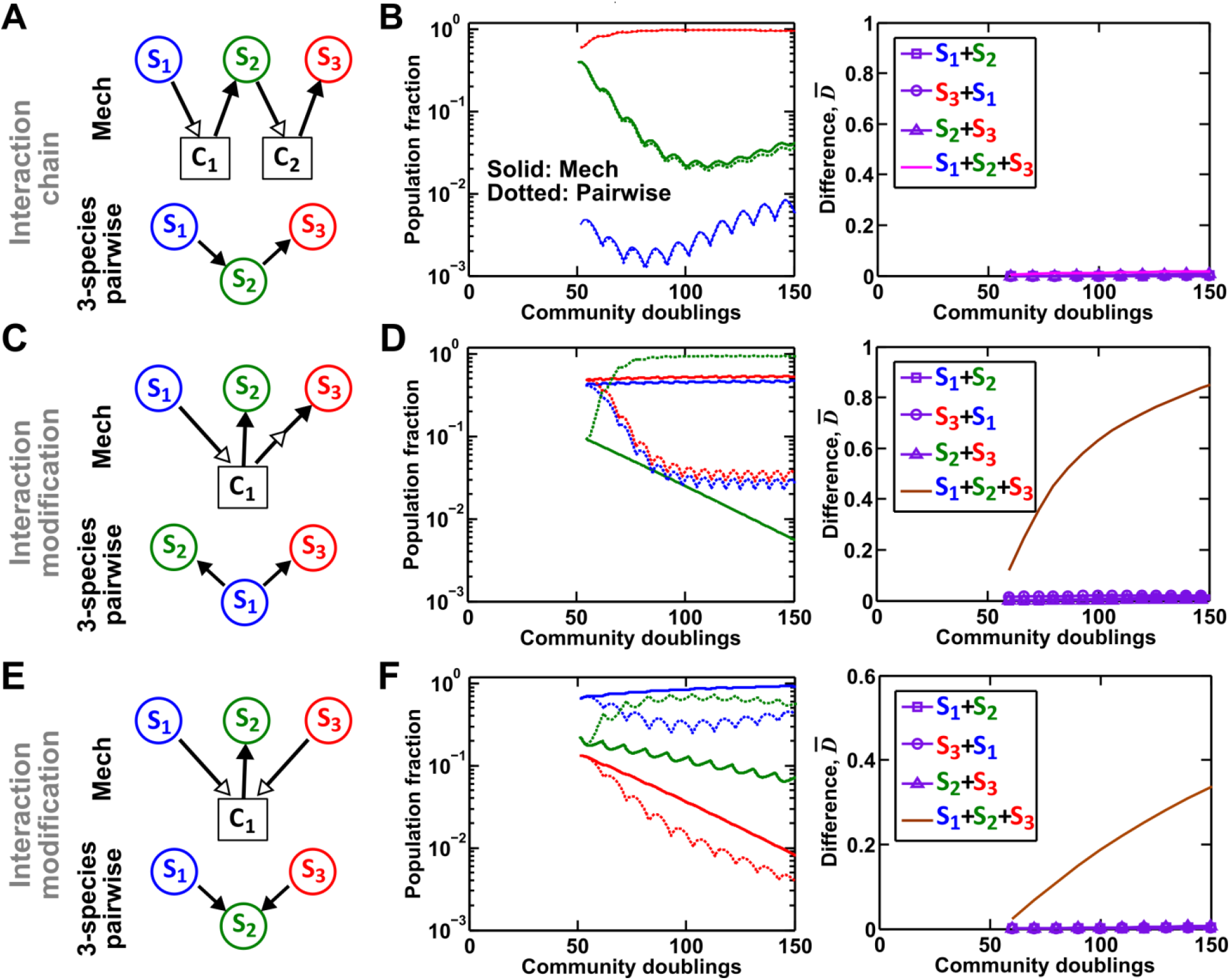
Interaction chain but not interaction modification may be represented by multispecies pairwise model. We examine three-species communities engaging in indirect interactions. Each species pair is representable by a two-species pairwise model (purple in the right columns of **B**, **D**, and **F**). We then use these two-species pairwise models to construct a three-species pairwise model, and test how well it predicts the dynamics from mechanistic model. In **B**, **D**, and **F**, left panels show dynamics from the mechanistic models (solid lines) and three-species pairwise models (dotted lines). Right panels show the difference metric *D* calculated over population densities after taking dilution into consideration. (**A**-**B**) Interaction chain: **S_1_** affects **S_2_**, and **S_2_** affects **S_3_**. The two interactions employ independent mediators **C_1_** and **C_2_**, and both interactions can be represented by the canonical pairwise model. The three-species pairwise model matches the mechanistic model in this case. Simulation parameters are provided in Fig 5-SD1. (**C-F**) Interaction modification. (**C-D**) **S_3_** consumes **C_1_**, a mediator by which **S_1_** stimulates S2. Parameters are listed in Fig 5-SD2. (**E-F**) **S_1_** and **S_3_** both supply **C_1_** which stimulates **S_2_**. Simulation parameters are listed in Fig 5-SD3. In both interaction modification cases, the three-species pairwise model fails to predict reference dynamics.

**Fig 6.**
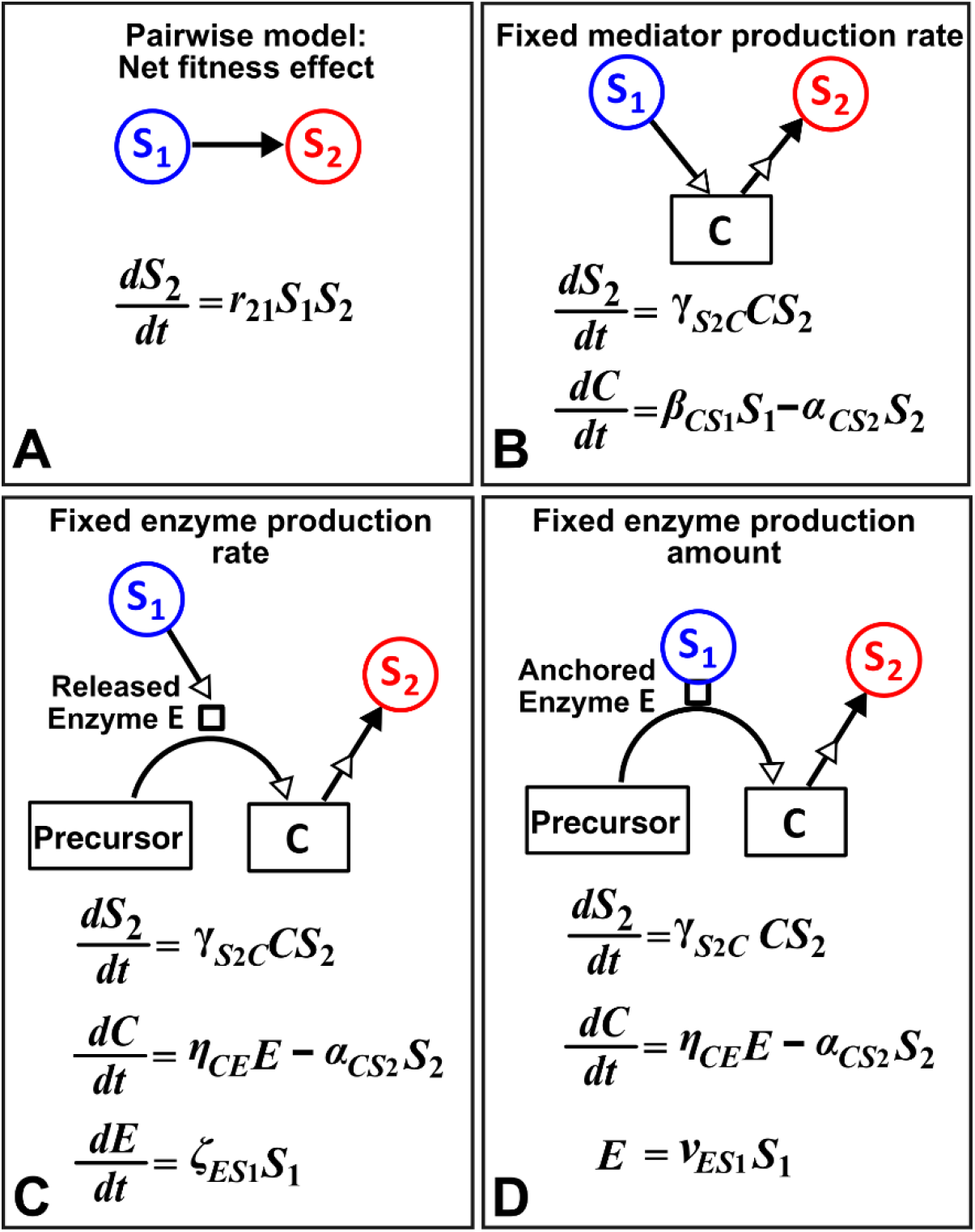
Different levels of abstraction in mechanistic modeling. How one species (**S_1_**) may influence another (**S_2_**) can be mechanistically modeled at different levels of abstraction. For simplicity, here we assume that interaction strength scales in a linear (instead of saturable) fashion with respect to mediator concentration or species density. The basal fitness of **S_2_** is zero. (**A**) In the simplest form, **S_1_** stimulates **S_2_** in a pairwise model. (**B**) In a mechanistic model, we may realize that **S_1_** stimulates **S_2_** via a mediator C which is consumed by **S_2_**. The corresponding mechanistic model is given. (**C**) Upon probing more deeply, it may become clear that **S_1_** stimulates **S_2_** via an enzyme **E**, where **E** degrades an abundant precursor (such as cellulose) to generate mediator **C** (such as glucose). In the corresponding mechanistic model, we may assume that **E** is released by **S_1_** at a rate ***ζ_ESX_*** and that **E** liberates **C** at a rate ***η*_*CE*_**. (**D**) If instead **E** is anchored on the cell surface (e.g. in cellulose degradation via cellulosome), then *E* is proportional to S1. If we substitute *E* into the second equation, then (**B**) and (**D**) become equivalent. Thus, when enzyme is anchored on cell surface but not when enzyme is released, the mechanistic knowledge of enzyme can be neglected.

## Supplementary figures

**Fig 1-FS1.**
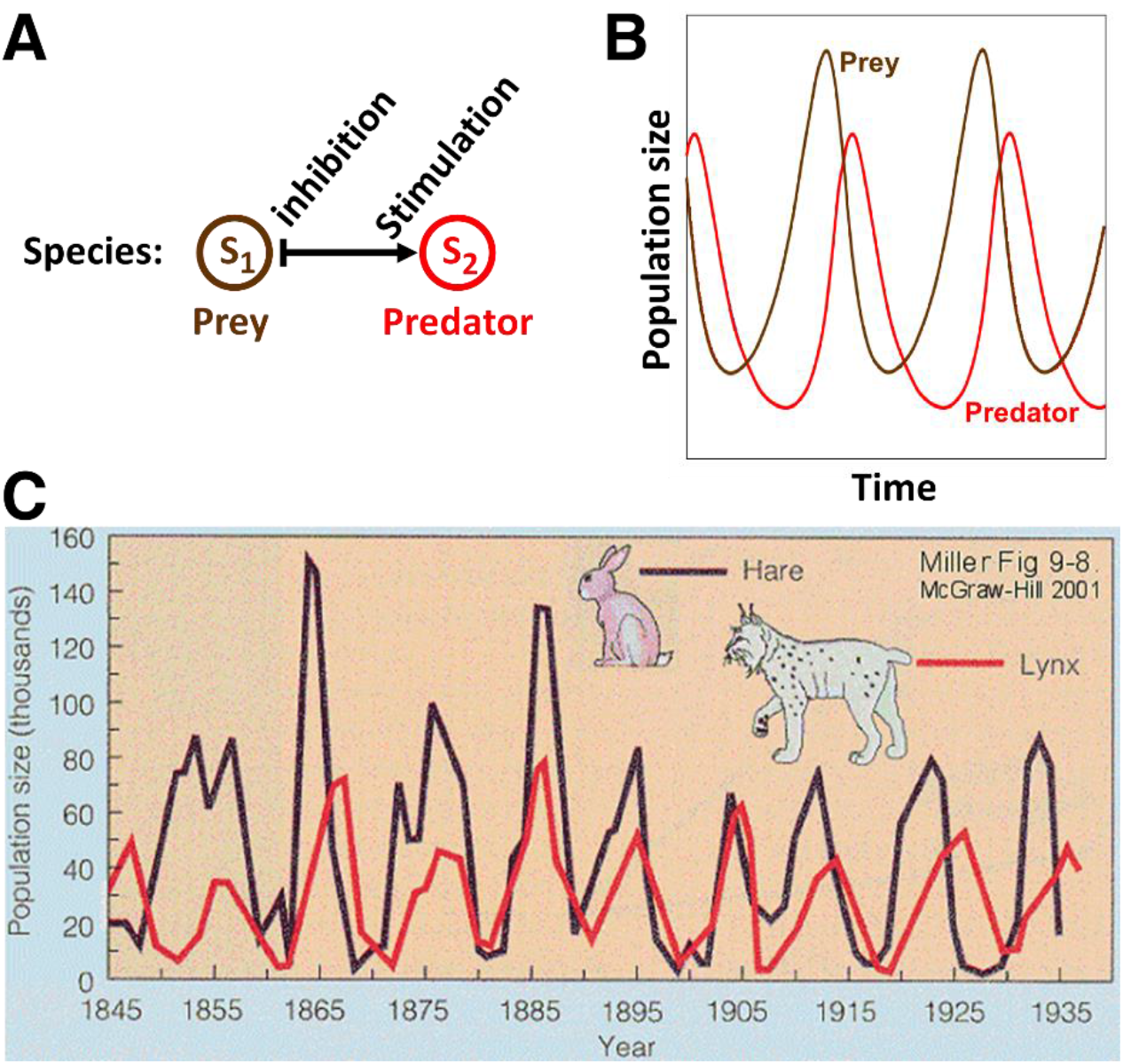
A pairwise model successfully predicts oscillations in population dynamics of the hare-lynx prey-predator community. (**A**) In a pairwise model of prey-predation proposed by Lotka and Volterra, predator reduces the fitness of prey, while prey stimulates the fitness of predator. (**B**) Assuming random encounter between prey and predator, the pairwise model predicts oscillations in the prey and predator population sizes. (**C**) Similar oscillations have been qualitatively observed in natural populations of lynx and hare, providing support for the usefulness of pairwise modeling. Picture is reproduced from (“BiologyEOC - PopulationChanges” 2016).

**Fig 1-FS2.**
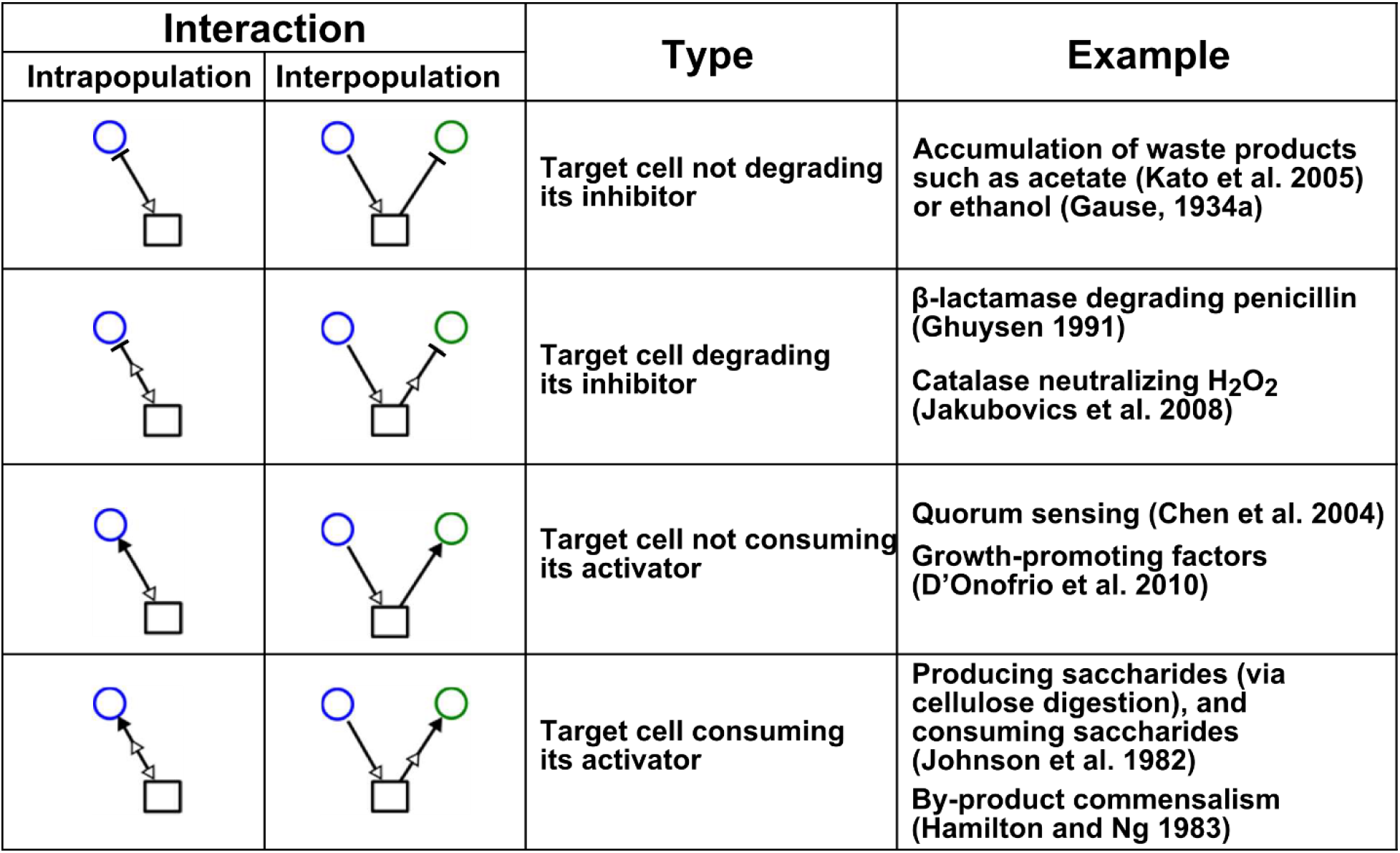
Chemical-mediated interactions commonly found in microbial communities. Interactions can be intra- or inter-population. Examples are meant to be illustrative instead of comprehensive.

**Fig 3-FS1.**
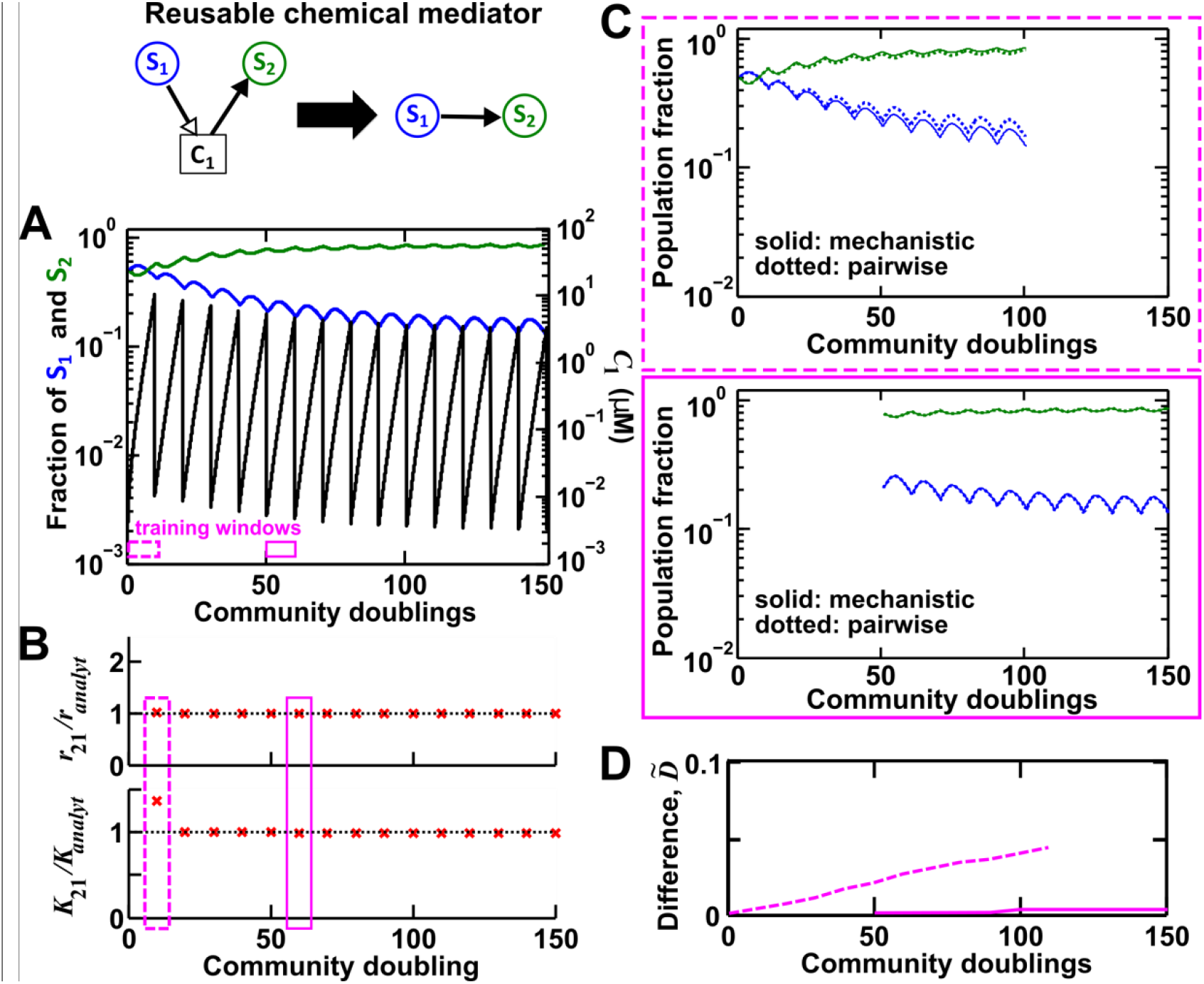
For a reusable mediator, parameter estimation after acclimation time leads to a more accurate canonical pairwise model. (**A**) We use the mechanistic model for a reusable mediator to generate reference dynamics of *Si, **S_2_***, and ***C_1_*** over 150 generations of community growth. The basal fitness of **S_1_** and **S_2_** in pairwise models are identical to those in mechanistic models, and here ***r_ii_*** or ***K_ii_*** (*i* = 1, 2) are irrelevant due to the lack of intrapopulation interactions. We use every 10 community doublings of reference dynamics as training windows to numerically estimate best-matching canonical pairwise model parameters ***r_21_*** and ***K_21_***. Dashed and solid rectangles represent training windows before and after acclimation, respectively. Note that population fractions (instead of population densities) are plotted, which fluctuate less during dilutions compared to mediator concentration. (**B**) Pairwise model parameters estimated after acclimation (solid rectangle) match their analytically-derived counterparts (black dotted lines) better than those estimated before acclimation (dashed rectangle). (**C**) A pairwise model generated from population dynamics before acclimation (top) predicts future reference dynamics less accurately than that generated after acclimation (bottom). (**D**) Quantification of the difference between pairwise and mechanistic models before (**dashed**) or after (**solid**) acclimation. All parameters are listed in Fig 3-SD1.

**Fig 3-FS2.**
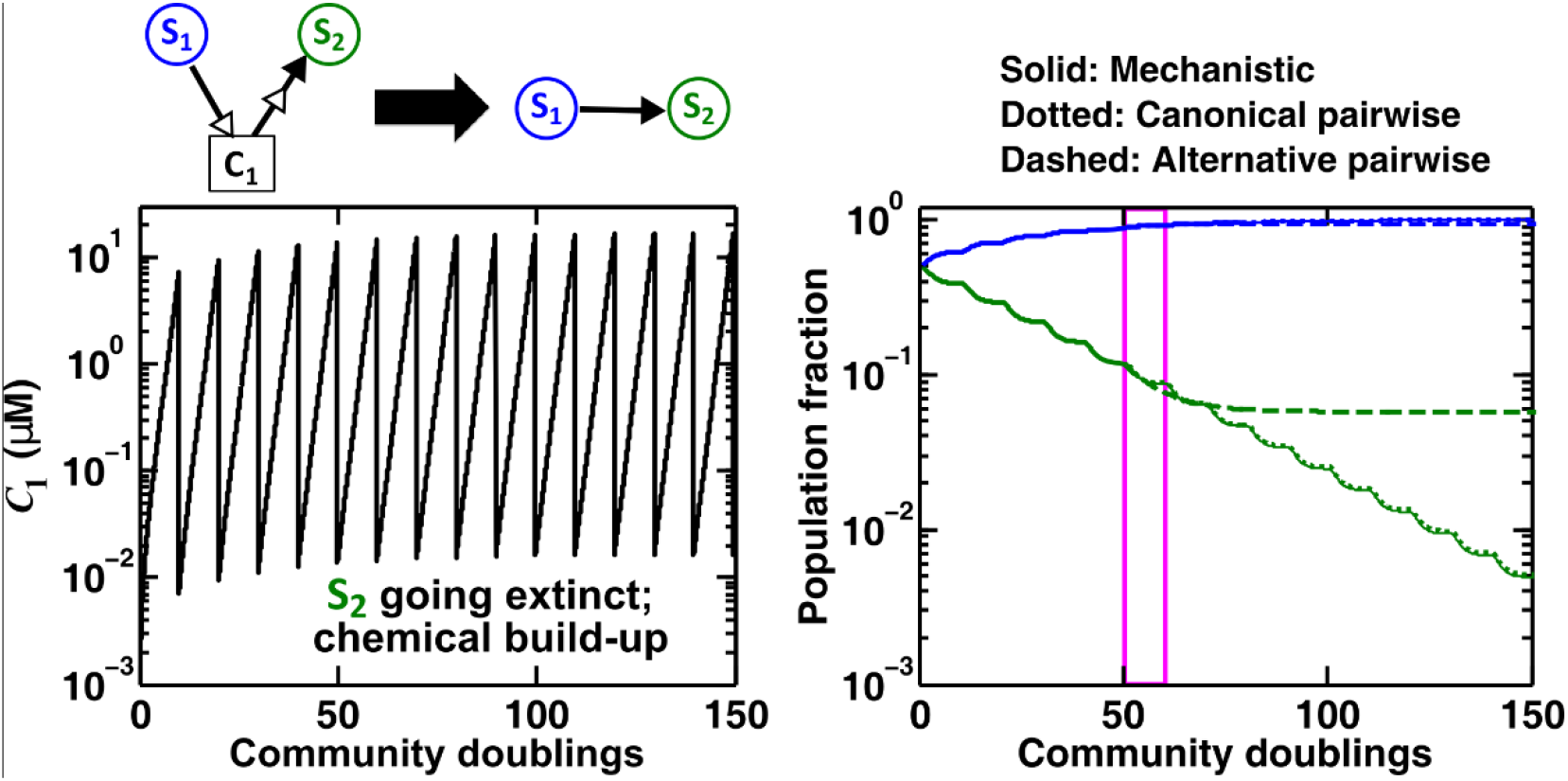
A canonical pairwise model, but not the alternative pairwise model, is suitable for a consumable mediator that accumulates without reaching a steady state within each dilution cycle. In a commensal community, the consumable mediator **C_1_** accumulates as the consumer **S_2_** gradually goes extinct. Pairwise model parameters are estimated from the mechanistic model dynamics in the training window (magenta, between 50 and 60 generations). The canonical model (Fig 3A) shows dynamics (dotted) that match those of the mechanistic model (solid). As expected, the alternative pairwise model (dashed) fails. Thus, accumulating **C_1_** can be regarded as a reusable mediator. Note that population fractions (instead of population densities) are plotted, which fluctuate less during dilutions compared to mediator concentration. All parameters are listed in Fig 3-SD2.

**Fig 3-FS3.**
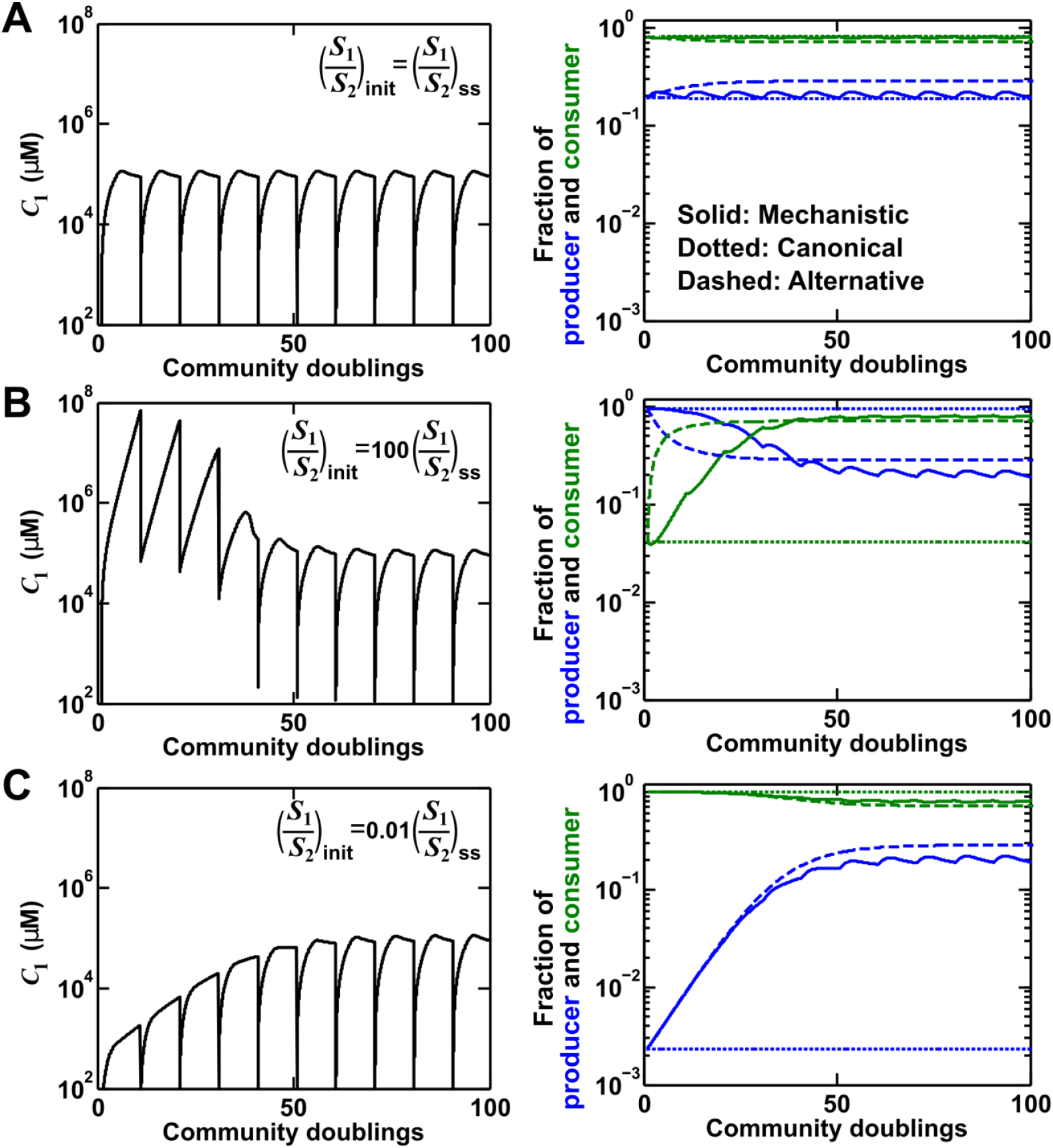
Interactions mediated by a consumable chemical that reaches a steady state can be represented by the alternative but not canonical pairwise model. Consider a commensal interaction where the consumable mediator reaches a non-zero steady state (Fig 3B, Case II). We can directly compute from the mechanistic model the corresponding canonical pairwise model (similar to Fig 3B, ii): 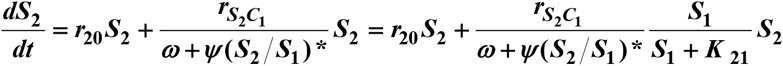 where ***K_21_*** < < ***S_1_***, and 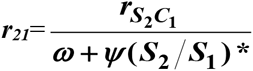. (**A**) As expected, when the community starts at the steady state, both the canonical and the alternative pairwise models predict steady-state dynamics. (**B** and **C**) When the community does not start at the steady state, the canonical model falsely predicts the maintenance of initial ratios. The alternative model predicts a convergence to the steady state, similar to the mechanistic reference model. Note that population fractions (instead of population densities) are plotted, which fluctuate less during dilutions compared to mediator concentration. All parameters are listed in Fig 3-SD3.

**Fig 3-FS4.**
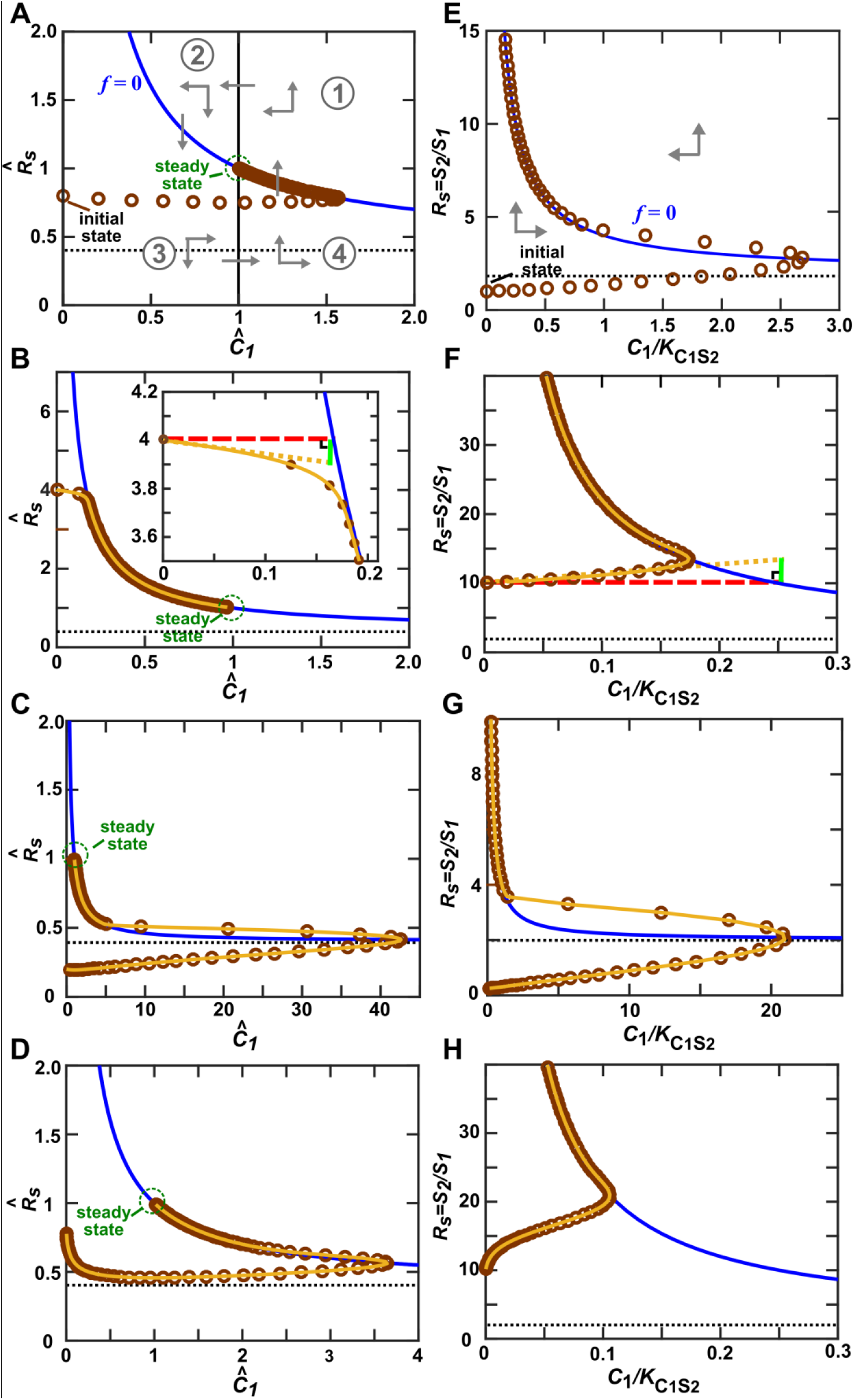
Community trajectory approaching the *f*-zero-isocline allows us to use the alternative pairwise model approximation. **S_1_** releases a consumable metabolite **C_1_** which stimulates **S_2_** growth. In all panels, brown circles indicate the **C_1_** and ***R_S_*** (=***S*_2_/*S*_1_**) of a community, and are separated by 1/4 of community doubling time. In the vicinity of the ***f***-zero-isocline (***f*** = 0) (blue line), ***C*_1_** can be eliminated to yield a pairwise model. (**A-D**) When ***r*_*S*_2_*C*_1__ > *r*_20_ − *r*_20_ > 0** (Methods-Section 3, Case II), a steady state (green circle) exists. Let us scale ***R_S_*** and **C_1_** against their respective steady state values to obtain 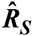 and 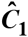. The ***f***-zero-isocline and the steady state 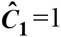 (vertical solid line) divide the phase portrait into four regions ((①) to (④)) (**A**). The directions of movement are marked by grey arrowheads. In Eq. S3-7, the right-hand side is zero when 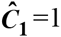. Since 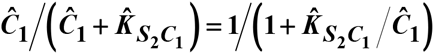 is an increasing function of 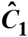, when 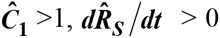 (up arrows), and when 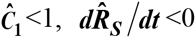 (down arrows). From Eq. S3-8, above the ***f***-zero-isocline, 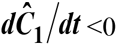 (left arrows), while below the f-zero-isocline, 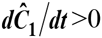 (right arrows). Thus, the community moves toward the f-zero-isocline, and then moves slowly alongside (but not superimposing) the ***f***-zero-isocline before reaching the steady state. **A-C** respectively describe community dynamics trajectories from *C*_1_ =0 when ***S*_10_** is large and when 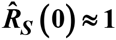 (Case II-2), 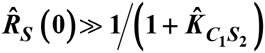 (Case II-1), or 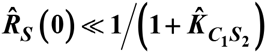 (CaseII-3). 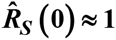 in D but ***S_10_*** is much smaller than that in A. In this case, instead of approaching the ***f***-zero-isocline quickly as in A, the trajectory plunge sharply before moving toward the ***f***-zero-isocline. The black dotted line marks 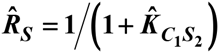, the asymptotic value off-zero-isocline. (**E-H**) When ***r*_10_ − *r*_20_** < 0 (Methods-Section 3, Case III), there is no steady state. ***R_S_*** approaches infinity and **C_1_** approaches 0. The black dotted line marks 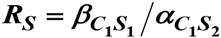, the asymptotic value of ***f***-zero-isocline. **E**, **F**, and **G** respectively describe community dynamics trajectories from **C_1_**= 0 when ***S*_10_** is large and when 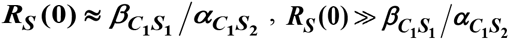, (Case III-1), and 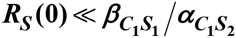 (Case III-2). In D, 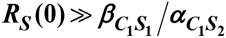 but ***S*_10_** is much smaller than that in F. Note different axis scales in different figure panels. All parameters are listed in Fig 3-SD4.

**Fig 3-FS5.**
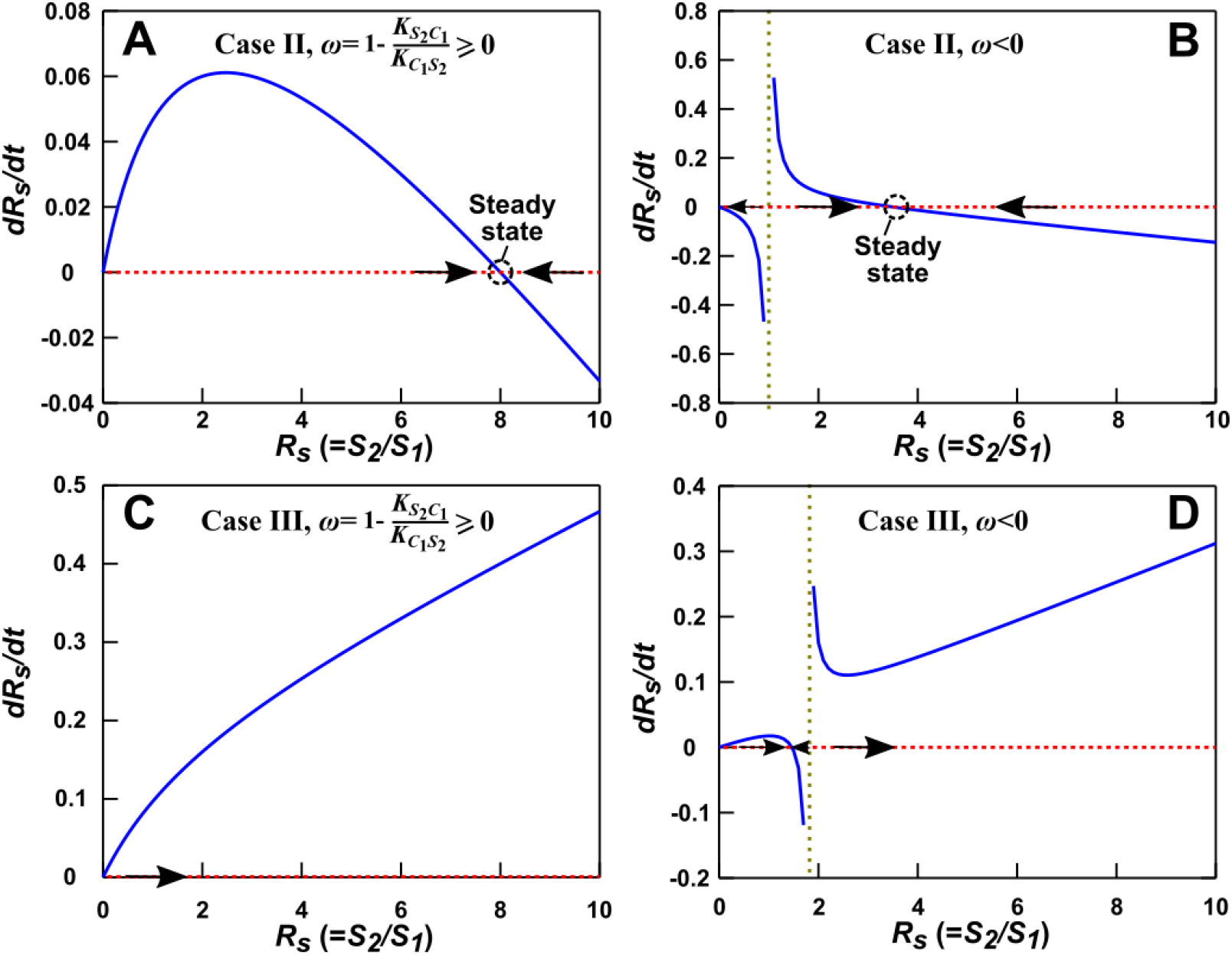
Condition for the alternative pairwise model to converge to mechanistic model. Here are the phase portraits of Eq. S3-31. The olive vertical dotted lines correspond to ***R_S_* = −ω/ψ**, a singularity point when ***ω*** < 0. (**A**) Case II (***r*_*S*_2_*C*_1__ > *r*_10_ − *r*_20_ > 0**), ***ω*** = 0.5, ***Ψ*** = 0.25. Regardless of initial ***R_S_***, the solution converges to steady state (in agreement with the mechanistic model). (**B**) Case II, ***ω*** = −1, *Ψ* = 1. When ***R_S_***(***t* = 0**) < −***ω/ψ*** (to the left of olive line), pairwise model falsely predicts extinction of ***S_2_***. (**C**) Case III (***r*_10_ < *r*_20_**), ***ω*** = 0.8, ***Ψ*** = 0.1. Regardless of initial ***R_S_***, the model predicts extinction of **S_1_** (in agreement with the mechanistic model). (**D**) Case III (***r*_10_ < *r*_20_**), ***ω*** = −9, ***Ψ*** = 5. When ***R_S_*** (***t* = 0)<−*ω/ψ*** (to the left of olive line), pairwise model falsely predicts steady state coexistence of the two species. All parameters are listed in Fig 3-SD5.

**Fig 3-FS6.**
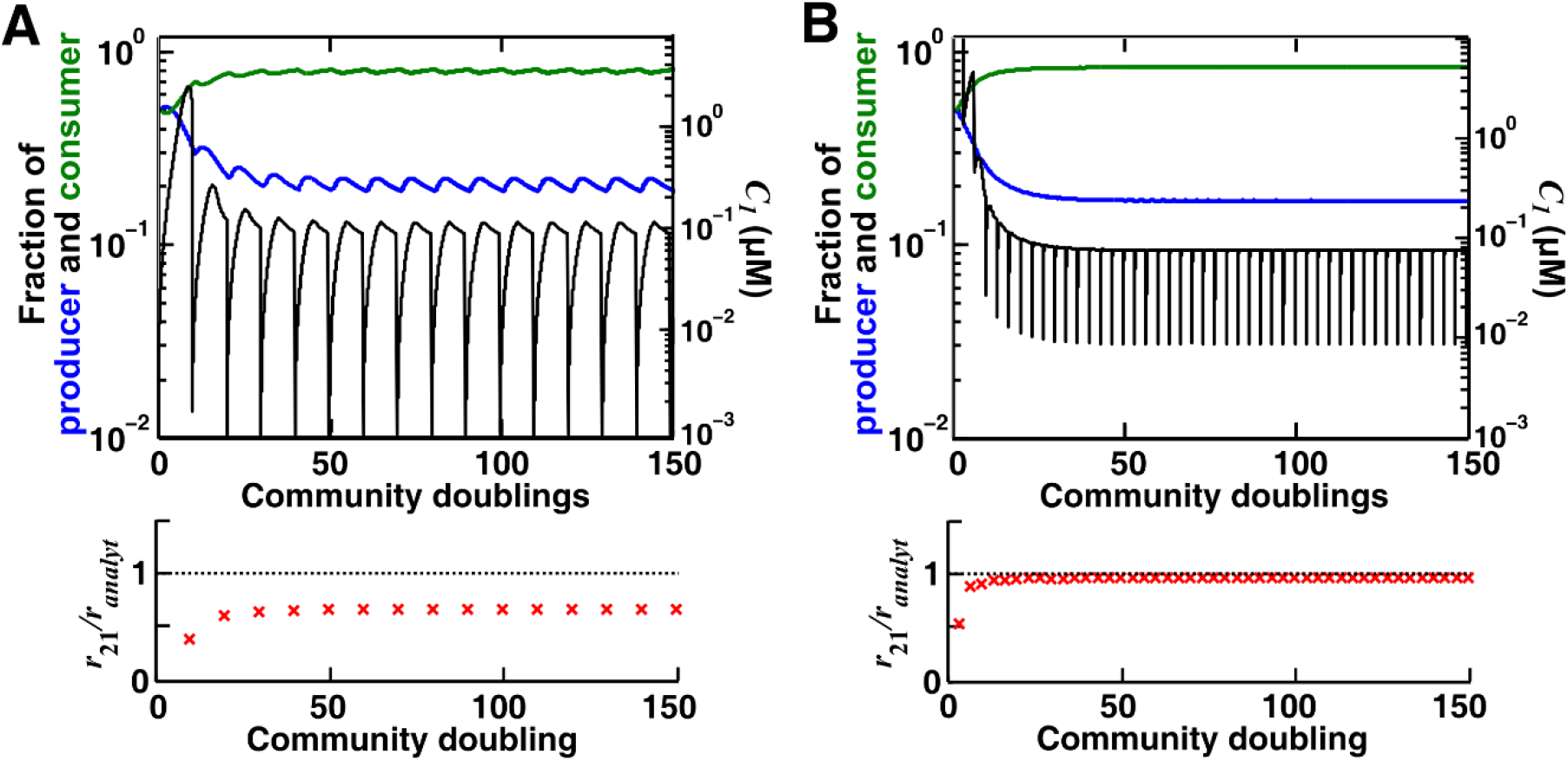
The accuracy of parameter estimation for the alternative pairwise model is influenced by time of estimation and dilution schemes. We consider commensalism through a consumable mediator, where the producer (blue) and the consumer (green) eventually reach a steady state. We compare 1000-fold (**A**) and 10-fold (**B**) dilution steps to examine how fluctuations caused by dilutions affect parameter estimation. We use every 10 community doublings of reference dynamics as training windows to numerically estimate best-matching alternative pairwise model parameters. In (**A**), compared to (**B**), we see larger errors in estimating the interaction strength ***r_21_*** compared to the true value (calculated from Fig 3B). In both (**A**) and (**B**), parameter estimations are less accurate if estimated before ***C*_1_** has stabilized (rigorously speaking, before the community dynamics has approached the ***f***-zero-isocline). Note that population fractions (instead of population densities) are plotted, which fluctuate less during dilutions compared to mediator concentration. All parameters are listed in Fig 3-SD6.

**Fig 3-FS7.**
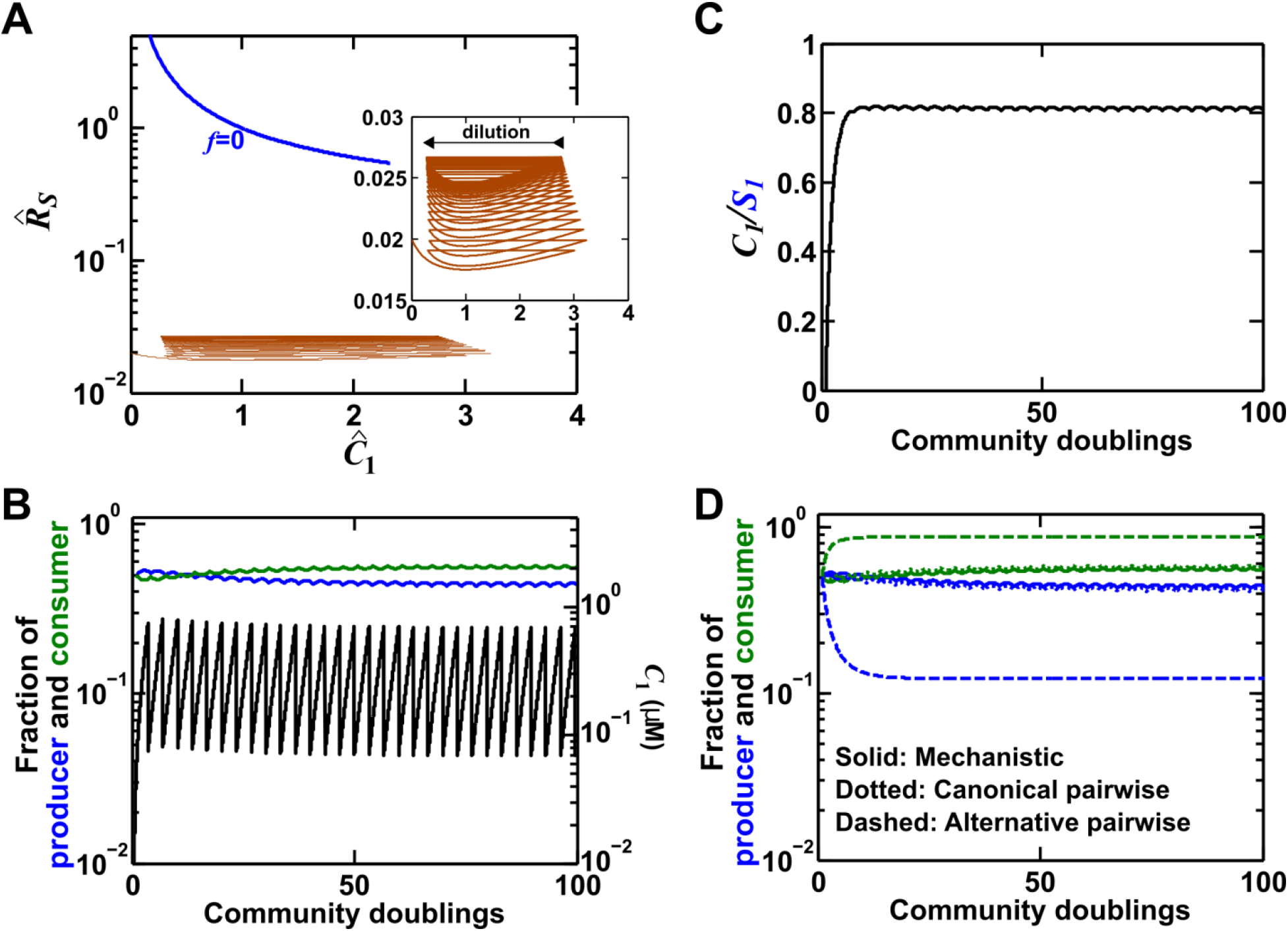
Dilutions may prevent a community from reaching the ***f***-zero-isocline required for the alternative pairwise model. We consider commensalism through a consumable mediator, where the producer (blue) and the consumer (green) could reach a steady state (Methods-Section 3 Case II). We choose a low consumption rate such that starting from equal proportions of producers and consumers, consumption can be neglected. (A) If the dilution scheme is set in a way that it prevents the community from approaching the blue ***f***-zero-isocline (brown trajectory in the phase space of 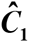 and 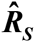, the mediator concentration and population ratio normalized to their potential steady state values, respectively), the community may reach an alternate sustained cycle. (**B**) In this case, the population fractions can remain steady (blue and green), while the mediator concentration appears to accumulate steadily within each dilution cycle without reaching a steady state (black). (**C**) In fact, ***C_1_*** appears to accumulate proportional to ***S_1_***. (**D**) Since the community remains far from the ***f***-zero-isocline, the use of the alternative pairwise model is not justified in this case. Instead, since the mediator concentration is proportional to the producer population size (**C**), the canonical pairwise model provides a better approximation. All parameters are listed in Fig 3-SD7.

**Fig 3-FS8.**
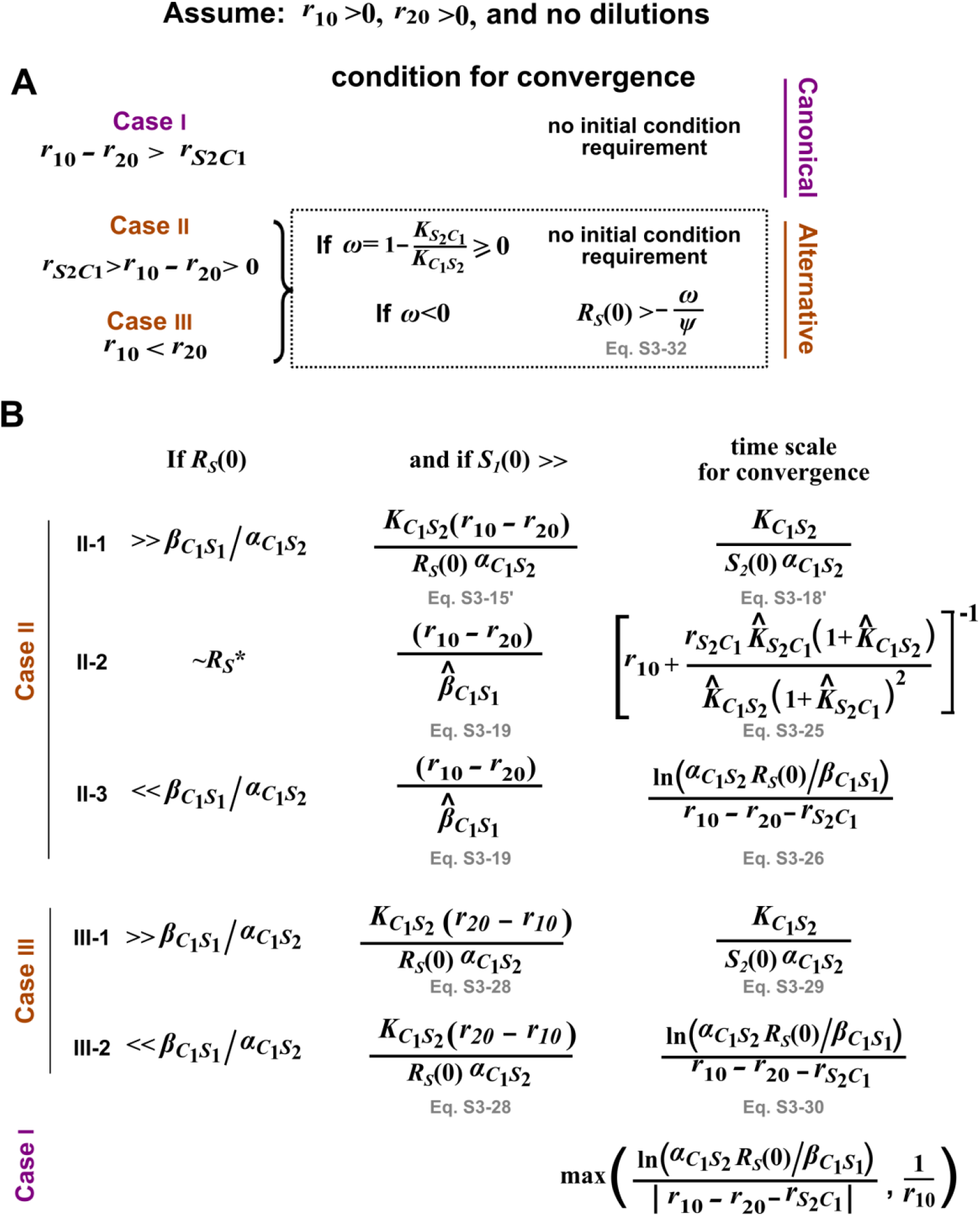
Additional requirements for deriving a pairwise model from mechanistic model when S_1_ affects S_2_ via a single consumable mediator C_1_ where ***C*_1_ (0) = 0**. For details, see Methods-Section 3. Here, ***R_S_*** = ***S*_2_/*S*_1_**. ***S***_10_, ***S***_20_, ***C*_1_ (0)**, and ***R_S_* (0)** are the initial values of the respective variables. (**A**) The initial condition requirement for a pairwise model to converge to the mechanistic model. (**B**) The time scale required for convergence. Conditions on *S*10 are sufficient, but may not be necessary.

**Fig 5-FS1.**
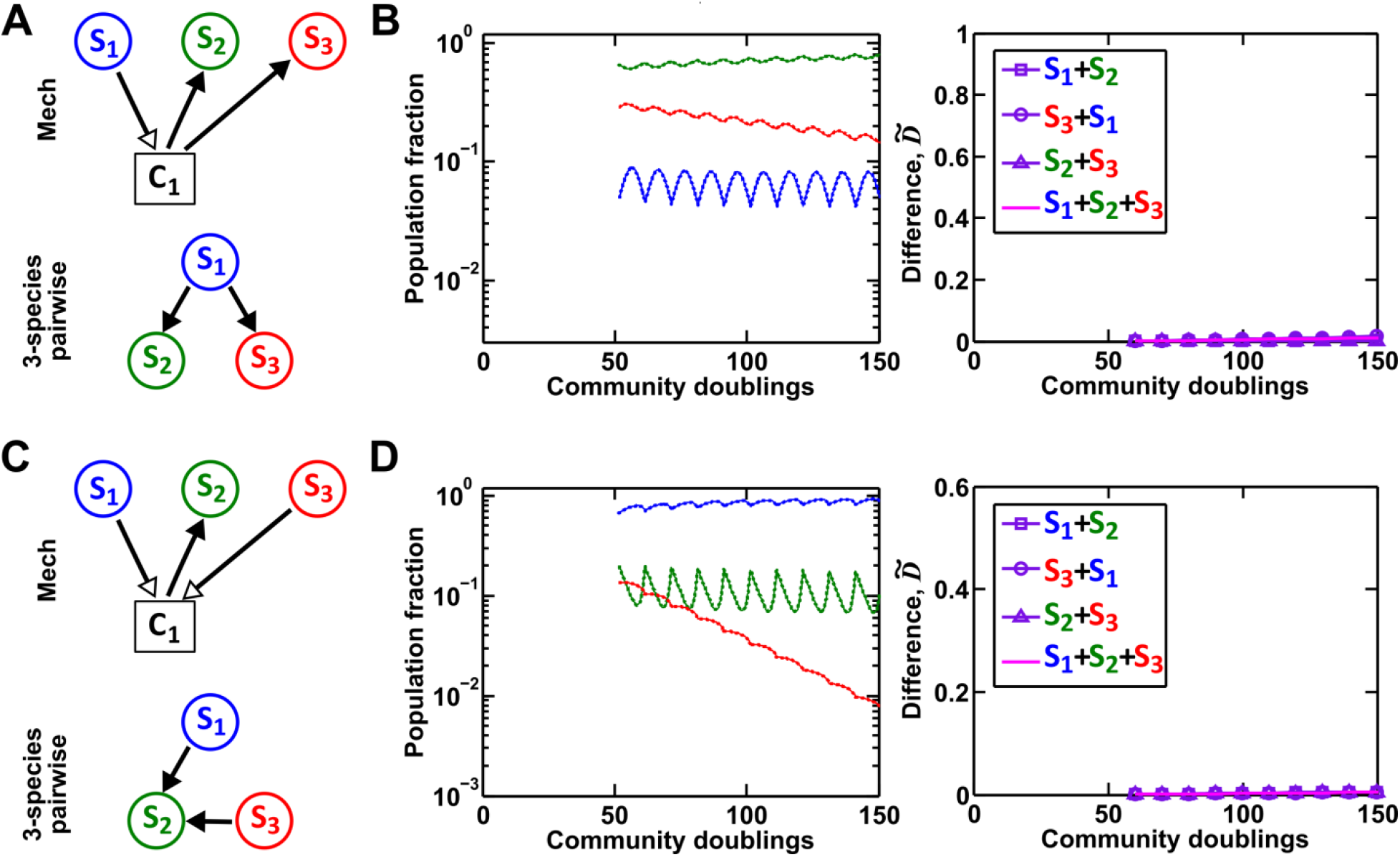
A multispecies pairwise model can work under special conditions. (**A-B**) As a control for Fig 5C, if **S_3_** does not remove the mediator of interaction between **S_1_** and **S_2_**, a three-species pairwise model accurately matches the mechanistic model. Simulation parameters are provided in Fig 5-SD4. (**C-D**) As a control for Fig 5E, we ensured that fitness effects from multiple species are additive. In this case, a three-species pairwise model can represent the mechanistic model. To ensure the linearity of fitness effects, we have used a larger value of half saturation concentration (***K*_*S*_2_*C*_1__** = 10^9^ instead of 10^5^ in Fig 5E-F). We have adjusted the interaction coefficient accordingly such that the overall interaction strength exerted by **S_1_** and **S_3_** on **S_2_** is comparable to that in Fig 5E-F (as evident by comparable population compositions). Since the interaction influences under these conditions remain in the linear range, the three-species pairwise model accurately predicts the reference dynamics. Simulation parameters are provided in Fig 5-SD5.

